# Identification of Novel Modulators of the ALT Pathway Through a Native FISH-Based Optical Screen

**DOI:** 10.1101/2024.11.15.623791

**Authors:** Benura Azeroglu, Simran Khurana, Shih-Chun Wang, Gianna M. Tricola, Shalu Sharma, Camille Jubelin, Ylenia Cortolezzis, Gianluca Pegoraro, Kyle M. Miller, Travis H. Stracker, Eros Lazzerini Denchi

## Abstract

A significant portion of human cancers utilize a recombination-based pathway, Alternative Lengthening of Telomeres (ALT), to extend telomeres. To gain further insights into this pathway, we developed a high-throughput imaging-based screen named TAILS (Telomeric ALT *In situ* Localization Screen), to identify genes that either promote or inhibit ALT activity. Screening over 1000 genes implicated in DNA transactions, TAILS revealed both well-established and novel ALT modulators. We have identified new factors that promote ALT, such as the nucleosome-remodeling factor CHD4 and the chromatin reader SGF29, as well as factors that suppress ALT, including the RNA helicases DDX39A/B, the replication factor TIMELESS, and components of the chromatin assembly factor CAF1. Our data indicate that defects in histone deposition significantly contribute to ALT-associated phenotypes. Based on these findings, we demonstrate that pharmacological treatments can be employed to either exacerbate or suppress ALT-associated phenotypes.

## Introduction

The progressive loss of telomeric repeats during DNA replication leads to telomere shortening and represents a barrier to unlimited proliferation^1–4^. To overcome this tumor suppressive barrier, cancer cells activate either telomerase to synthesize new telomeric repeats or, in approximately 10-15% of all cancers, engage the Alternative Lengthening of Telomeres (ALT) pathway. The ALT pathway utilizes break-induced replication (BIR) to elongate telomeres, allowing telomeric DNA synthesis by using another telomere as a template^5, 6^. Major advances in understanding the ALT pathway have been made in recent years (reviewed in^7–9^). Nevertheless, several key aspects of the ALT pathway remain unclear, including a comprehensive analysis of cellular factors and pathways that both suppress and promote ALT activity. With the goal of identifying such factors, we set out to perform a high-throughput screen to identify potential novel modulators of the ALT pathway.

ALT telomeres display unique characteristics, including clustering within structures known as ALT-associated Promyelocytic Leukemia (PML) bodies (APBs) that contain several DNA damage response (DDR) proteins^10^. ALT-positive cells exhibit replication stress, increased levels of DNA-RNA hybrids and elevated DNA damage signaling at telomeres^11, 12^. Loss of ATRX or DAXX, histone chaperones that deposit the histone variant H3.3 in repetitive DNA, are a hallmark of ALT-positive cells^13–16^. As a result, ALT cells show a reduced deposition of H3.3 at telomeres, leading to elevated expression of telomeric repeat-containing RNA (TERRA)^11^. TERRA-associated R-loops are stabilized by G-quadruplex formation and by the RAD51AP1 protein^17, 18^.

Among the proteins required to sustain ALT-mediated telomere elongation in cancer cells, the Bloom syndrome DNA helicase BLM is particularly important. BLM is part of the BLM-TOP3A-RMI1 (BTR) complex, which is recruited to telomeres by PML and is required to sustain ALT activity^19^. BLM helicase activity plays a key role in the assembly of DNA repair proteins on ALT telomeres, and it has been shown to play a role in the processing of lagging strand intermediates in conjunction with the nuclease DNA2^20^.

There is considerable interest in developing therapies tailored for ALT-positive cancers, given that they would target a significant number of human cancers^21, 22^. Multiple therapeutic strategies have been proposed to selectively kill ALT-positive cells. Broad-spectrum approaches include small molecules targeting the DDR pathway, such as ATR or PARP1 inhibitors^23, 24^. More targeted approaches have been proposed, including the inhibition of FANCM, an ATPase/translocase enzyme that is part of the Fanconi Anemia DNA repair complex, or KDM2A, a histone demethylase^25, 26^. FANCM depletion induces toxic hyper-ALT phenotypes in ALT-positive cells and depletion of KDM2A was shown to prevent ALT-telomere declustering, leading to mitotic damage and cell death^26–28^. Additionally, the loss of specific genes, including HIRA, SMC5/6 family members, SMARCAL1, and WEE1, or the amplification of MYCN, were found to sensitize ATRX-deficient cells, suggesting additional interventions to selectively target ALT cells^29–33^. Finally, since ALT cells typically silence the innate immune cytoplasmic dsDNA sensor STING, reactivating STING has been proposed as an approach to trigger immunogenicity in ALT-positive tumors^34, 35^.

Here, we aimed to identify additional modulators of ALT to deepen our understanding of its regulation and genetic dependencies. To achieve this, we developed a high-throughput imaging-based screening assay based on the detection of telomeric single-stranded DNA (ssDNA) using the ssTelo assay that we previously established^19^. We and others have shown that the ssTelo assay can sensitively detect ssDNA at telomeres, which correlates with ALT activity^19, 36, 37^. This assay was combined with a loss-of-function CRISPR Knockout (KO) screen using a focused, arrayed sgRNA library.

Our screen identified 6 known and 85 novel putative modulators of ALT activity. We validated and characterized 7 of these novel genes as activators or suppressors of the ALT pathway, demonstrating that several of them affect histone deposition. These findings indicate that ALT-positive cells may be particularly vulnerable to disruptions in cellular processes reliant on histone deposition, such as DNA replication and transcription. Additionally, our screen identified several druggable proteins and pathways, providing a basis for designing drugs to either suppress or hyperactivate the ALT pathway. As a proof of concept, we identified several small-molecule inhibitors that can effectively modulate ALT-related phenotypes in human cancer cells.

## Results

### Telomeric ALT *In situ* Localization Screen (TAILS): a native-FISH based imaging screen

We previously reported that the presence of telomeric ssDNA could be used to discriminate ALT-positive cell lines from non-ALT cell lines^19^. Furthermore, we showed that telomeric native-FISH (ssTelo) could be used to assess changes in ALT activity within ALT-positive cell lines^19,38^. Based on this finding, we combined ssTelo staining with CRISPR-mediated gene deletion to perform a high-throughput functional genomics screen to identify modulators of the ALT process. We termed this approach TAILS for Telomeric ALT *In situ* Localization Screen.

To validate this approach, we performed TAILS in the ALT-positive osteosarcoma cell line U2OS using 3 pooled sgRNA oligos against non-targeting sequences (sgCtrl), the Bloom DNA helicase (sgBLM), and the DNA helicase FANCM (sgFANCM). The results of these experiments show that TAILS can detect a reduction in ALT activity following depletion of BLM, as well as an induction in ALT activity following FANCM loss, to a level that is comparable to the rolling circle assay (RCA), the current gold-standard assay for ALT activity (Figures 1A and 1B).

**Fig 1:**
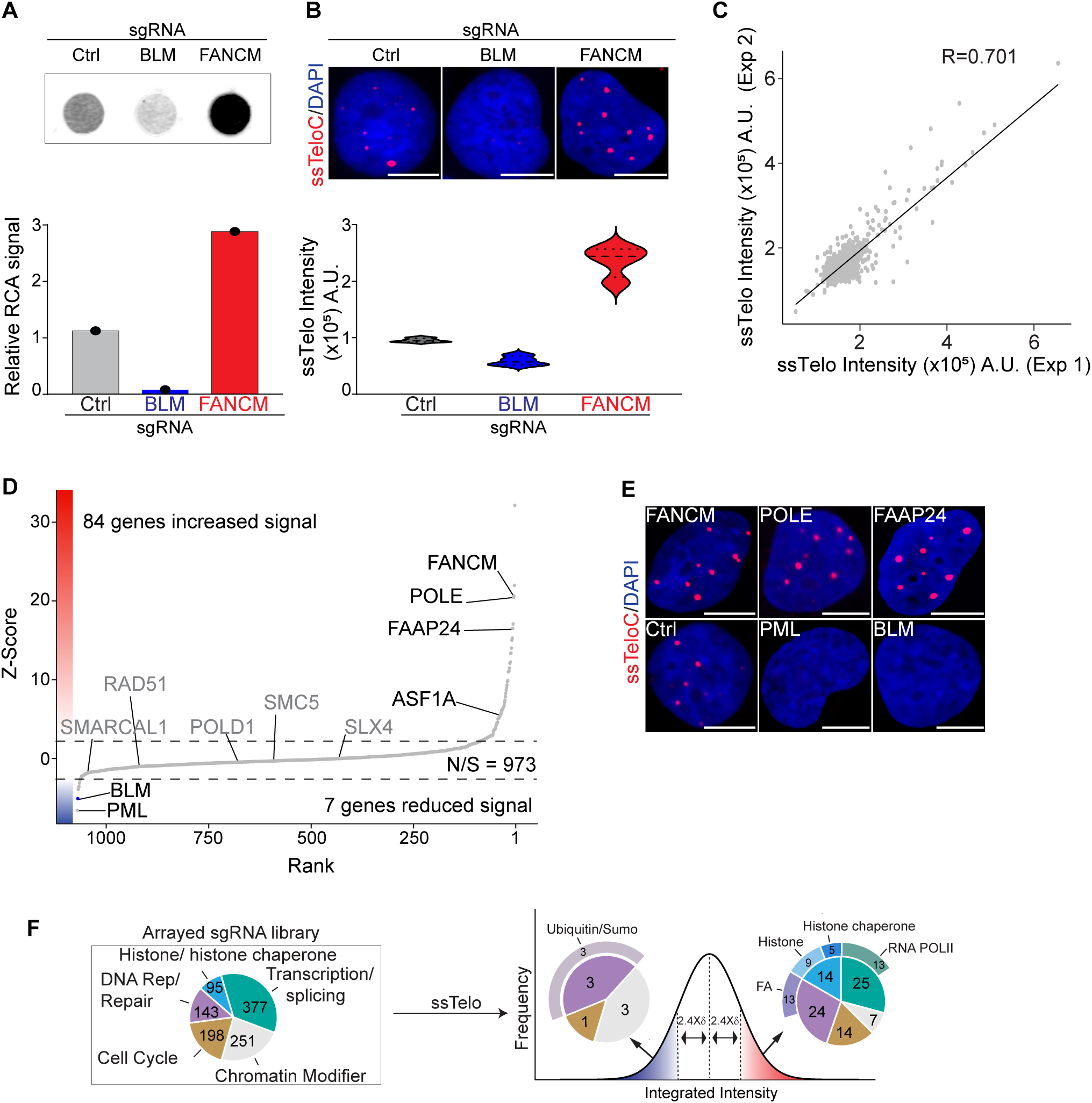
TAILS identifies novel modulators of ALT. **A** Rolling circle assay (RCA) analysis of genomic DNA isolated from U2OS cells transfected with the sgRNA and relative quantification. **B** Example of TAILS data and quantification; U2OS cells transfected with indicated sgRNA were processed as described in the TAILS pipeline. The scale bar is 10 μm. **C** TAILS analysis in ALT-positive U2OS cells. The ssTelo intensity derived from two biological replicates of the arrayed sgRNA library. Linear regression analysis by Pearson correlation coefficient (R) was calculated, and it is 0.701. **D** Scatter plot displaying mean Z-score for each gene assayed (y-axis) and the relative ranking based on descending Z-score (x-axis). Dotted lines indicate the cut-off value (+/-2.4) chosen to identify putative hits. Genes that have previously been reported to affect ALT activity are indicated. **E** Images obtained by TAILS of hits selected known modulators of ALT. The scale bar is 10 μm. **F** Gene function classification of all the genes contained in sgRNA library (left diagram) and of the hits identified by TAILS (right diagrams). For more information, see Supplementary Table 1.

Next, we performed TAILS using an arrayed sgRNA library consisting of 1064 genes implicated in DNA transactions (see Supplementary Table 1). 96 hours post-sgRNA transfection, cells were fixed, stained for ssTelo, and counterstained with DAPI (Figures S1A-C). Individual wells were scored for ssTelo foci number, ssTelo foci intensity, and ssTelo integrated intensity (Figures S1D-F). As controls, we included non-targeting sgRNAs (sgCtrl), gRNAs targeting the essential gene PLK1 (sgPLK1), and gRNAs targeting BLM (sgBLM). These controls were used to establish a baseline for ssTelo signal (sgCtrl and sgBLM) and to control for the efficiency of CRISPR-KO (sgPLK1) (Figures S1A-F). We performed the screen in two biological replicates and found that the data obtained for sgRNA replicates was reproducible, with a Pearson correlation coefficient of 0.7 calculated in a linear regression analysis (Figure 1C). To evaluate the effect of CRISPR-KO depletion of each individual gene on ALT, we calculated a Z-score based on the deviation of ssTelo signal intensity from the median value for all the sgRNAs present in the library. Genes with a Z-score below −2.4 or above 2.4 were considered putative ALT-activators or ALT-suppressors, respectively. Following this criterion, cells treated with sgBLM guides all scored as hits in the screen, while none of the negative control samples (sgCtrl) did. In total, we identified 84 genes as putative ALT-suppressors (Z-score > 2.4) and 7 genes as putative ALT-activators (Z-score < −2.4) (Figure 1D).

Among the putative ALT-suppressors, we found FANCM and 4 additional genes that several groups have previously characterized (see Supplementary Table 2 and Figure 1E)^26–28^. Additionally, we identified 24 genes involved in DNA replication/repair, including 13 genes of the Fanconi Anemia complex, 14 genes encoding histones or histone chaperones, and 25 genes involved in transcription, including 13 components of the RNA polymerase II (RNAPII) complex (Figures 1F and S1G-J, and Supplementary Table 1).

Among the putative ALT-activators, in addition to PML and BLM that are required to sustain ALT activity, we found 3 genes involved in DNA replication and repair via SUMO-mediated signaling, 3 chromatin remodeling factors, as well as a phosphatase involved in cell cycle regulation (Figure 1F and Supplementary Table 1)^19^. Collectively, our data show that using TAILS, we identified known modulators of the ALT pathway, as well as several putative novel ALT modulators involved in different biological processes.

### ALT-activators: SGF29, CHD4 promote ALT activity at telomeres

In addition to PML, we found other components of PML bodies, including the small ubiquitin-like modifier 2 (SUMO2) and the SUMO ligase UBE2I/UBC9 that reduced ssTelo signal when depleted. Additionally, we identified the Protein Phosphatase 2A subunit (PPP2CA) and multiple components of chromatin modifying complexes, including CHD4, EP300, and SGF29 as ALT-activators (Figures 2A and 2B, and Supplementary Table 1).

**Fig 2:**
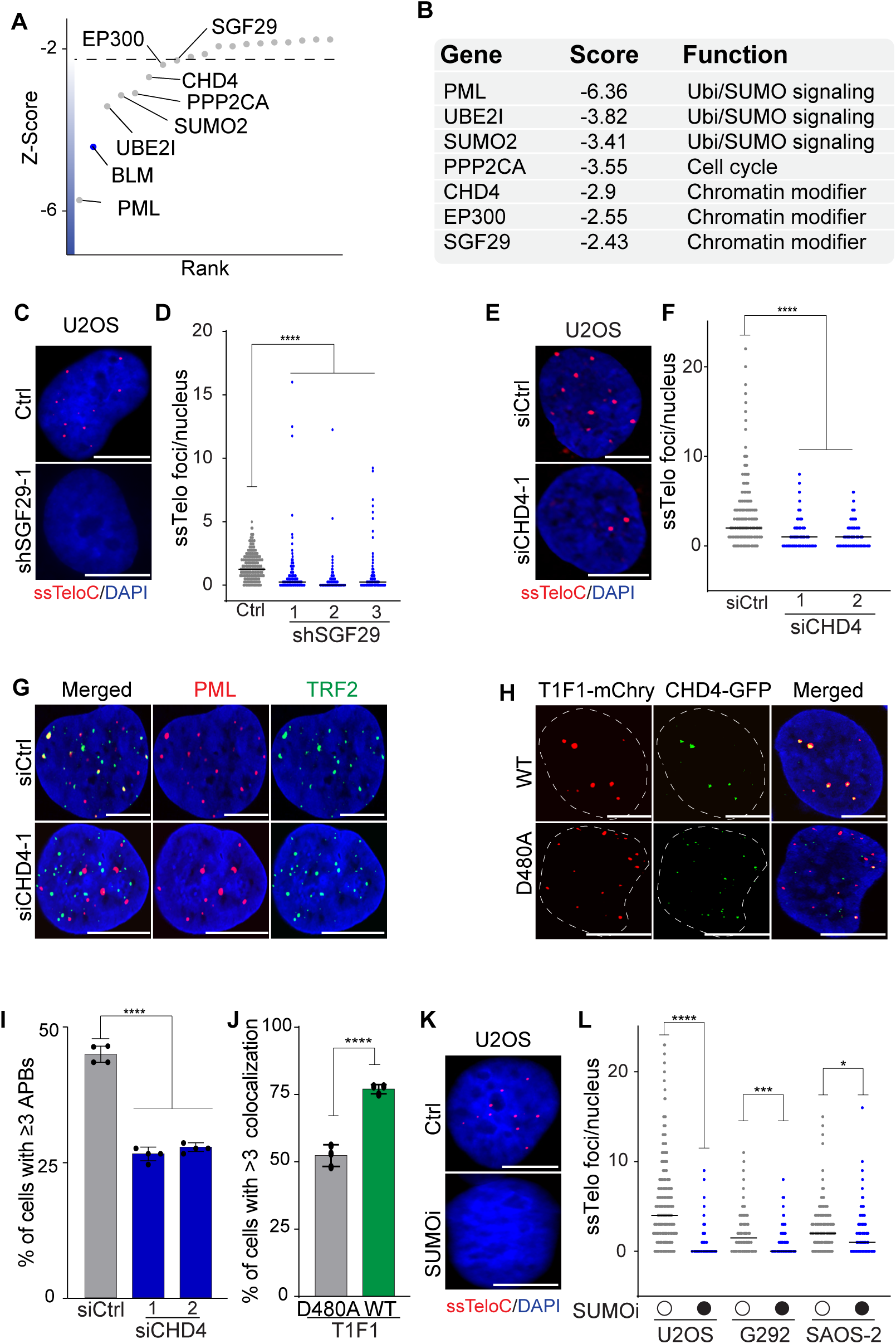
SGF29, CHD4 and SUMOylation are required for ALT activity. **A** Scatter plot shows the Z-score for genes identified by TAILS as putative ALT-activators. **B** Gene function classification of the genes displayed in panel A. **C-D** ssTelo staining and relative quantification of U2OS cells infected with shRNA against SGF29 (shSGF29-1, shSGF29-2, and shSGF29-3) or control (WT). **E-F** ssTelo staining and relative quantification of U2OS cells transfected with siRNA against CHD4 (siCHD4-1 and siCHD4-2) or a non-targeting control (siCtrl). **G** U2OS cells transfected with siRNA against CHD4 (siCHD4-1) or a non-targeting siRNA (siCtrl) were stained for PML (red) and TRF2 (green). **H** U2OS cells co-transfected with vectors expressed CHD4-GFP (green) and either WT or catalytically-dead (D480A) TRF1-FokI-mCherry (red). **I** Quantification of data shown in G with graphs indicating the percentage of cells with at least 3 PML-TRF2 colocalizations (APBs) per nucleus, defined as two foci overlapping by 50% or more. **J** Quantification of data shown in H with graphs indicating the percentage of cells with at least 3 CHD4-TRF1-FokI colocalizations per nucleus, defined as two foci overlapping by 50% or more. **K** ssTelo staining of ALT-positive U2OS, SAOS-2 and G292 cells treated with SUMOi compared to untreated sample. **L** Quantification of ssTelo analysis shown in K and S2H. The scale bar is 10 μm. An unpaired t-test was used for statistical analysis; P ≤ 0.05 indicated as *, P ≤ 0.001 indicated as ***, P ≤ 0.0001 indicated as **** on the graphs.

SGF29 is a chromatin reader and part of the histone acetyltransferase (HAT) module of both the SAGA and ATAC complexes involved in transcriptional regulation^39–41^. We depleted SGF29 using 3 independent shRNAs (Figure S2A) and found that SGF29 depletion significantly reduced the levels of ssDNA at telomeres (Figures 2C and 2D). This validated the data from the TAILS screen that suggested SGF29 was required to sustain ALT activity, however depletion of SGF29 did not significantly impair the proliferation of U2OS cells (Figure S2B).

Next, we examined the role of CHD4, an ATP-dependent nucleosome remodeler that is part of the NuRD histone deacetylase complex^42^. The NuRD complex is enriched in the chromatin of ALT telomeres, where it reduces telomeric histone acetylation and promotes ALT^43^. In addition, CHD4 has an established role in genome stability^44–47^. To test the role of CHD4 at telomeres, we performed siRNA-mediated gene silencing using two independent siRNAs. Our data showed that CHD4 depletion in U2OS cells resulted in a marked reduction in ssTelo signal as well as a reduction in the level of ALT-associated C-circles, as detected by the RCA assay (Figures 2E, 2F, and S2C-E). Similar results were obtained in the ALT-positive cell line LM216J (Figure S2G). We then tested whether the depletion of CHD4 affected the association of PML with telomeres, a critical step in the ALT process previously linked to NuRD complex recruitment^43^. We observed a marked decrease in the levels of ALT-associated PML bodies in CHD4 depleted cells, suggesting that the chromatin remodeling activity of CHD4 plays a pivotal role in recruiting PML to telomeres in ALT-positive cells (Figures 2G and 2I).

To further understand the role of CHD4 in the ALT process, we employed the TRF1-FokI endonuclease experimental system that generates breaks at telomeres and stimulates BIR-mediated ALT activity^48^. Expression of wild-type TRF1-FokI elevated CHD4 co-localization with telomeres compared to cells expressing a catalytically-dead allele of FokI (D480A), suggesting that CHD4 recruitment to telomeres might be promoted by DNA damage induction (Figures 2H and 2J). Together, these results suggest that CHD4 activity is required at telomeres to sustain the ALT process potentially by favoring the formation of APBs and the resulting recruitment of DNA repair factors involved in the ALT process. Depletion of CHD4 had a strong effect on the proliferation of U2OS cells (Figure S2G), a phenotype unlikely to be mediated by the loss of ALT activity itself, as suppression of ALT activity alone is insufficient to curb the proliferation of U2OS cells^19^.

Another pathway enriched among the ALT-activators was the SUMOylation pathway, represented by PML, SUMO2 and UBE2I (Figures 2A and 2B). SUMOylation is well-established to play a key role in ALT^49, 50^. To test whether ALT activity could be suppressed through pharmacological inhibition of the SUMO pathway, we treated cells with the SUMOylation inhibitor TAK-981 (referred to subsequently as SUMOi) at various doses and found that it had a marked effect on cell growth regardless of ALT status of the cells (EC_50_ in U2OS = 1.098 µM, EC_50_ in HeLa = 0.847 µM) (Figure S2I). Treatment with SUMOi suppressed the ssTelo signal in U2OS cells, as well in ALT-positive G292 and SAOS-2 cells, consistent with a role for this pathway in facilitating ALT (Figures 2K,2L, and 2SH). Together, these data show that our approach successfully identified both known and novel factors required for ALT activity.

### ALT-suppressors: a role for the transcription machinery and the RNA helicases DDX39A/B

We identified 84 putative ALT suppressors, of which 25 are involved in transcription and splicing, including most of the RNAPII subunits contained in the library (13 out of 17) (Figure 3A). To validate these findings, we first focused on DDX39A (DExD-box helicase 39A), a poorly characterized RNA helicase implicated in transcriptional gene regulation, shown to localize to ALT telomeres using proximity labeling and recently implicated in telomere maintenance in telomerase positive cells through interactions with TRF1^51, 52^.

**Fig 3:**
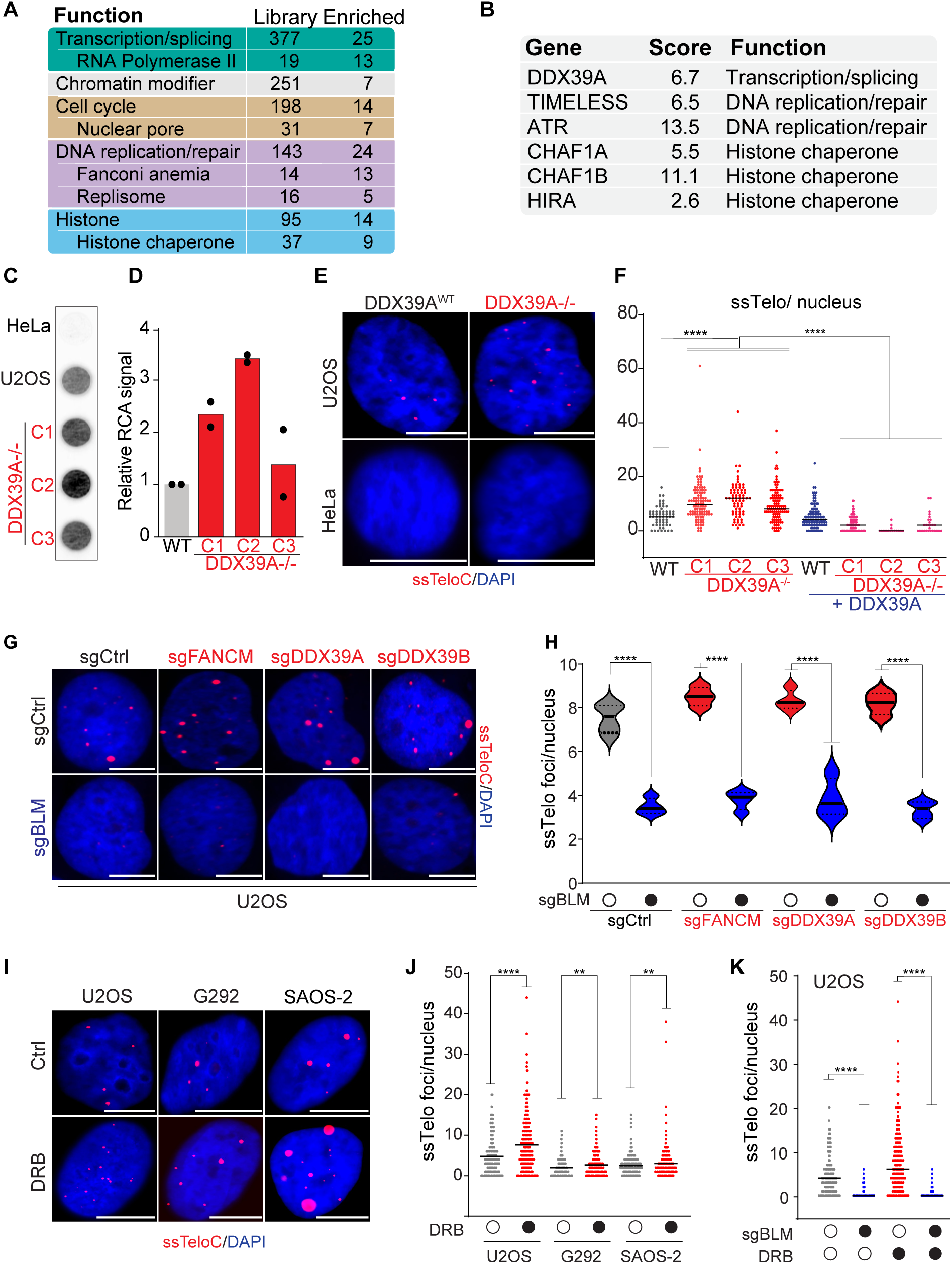
The RNA helicases DDX39A/B are ALT-suppressors. **A** Gene function composition of the arrayed sgRNA library (Library) and of the number genes that were identified as putative ALT suppressors by TAILS (Enriched). **B** List of ALT-suppressor genes that were further characterized in this study. The table reports the average Z-score (Score) and gene function category (Function). **C-D** Rolling circle assay (RCA) analysis of genomic DNA isolated from 3 independent DDX39A^-/-^ clones (C1, C2 and C3) and the parental U2OS cells (WT). **E-F** Representative images and quantification of DDX39A-deficient and - proficient cells in ALT-positive U2OS and ALT-negative HeLa backgrounds. The scale bar is 10 μm. **G** ssTelo staining of U2OS cells transfected with sgRNAs against FANCM, DDX39A, DDX39B and a non-targeting control (sgCtrl) in the presence of absence of sgRNA against BLM. The scale bar is 10 μm. **H** Quantification of the data shown in G. **I-J** Representative images and quantification of ssTelo analysis ALT-positive U2OS, G292 and SAOS-2 cells treated with transcription inhibitor DRB. The scale bar is 10 μm. **K** Quantification of ssTelo analysis U2OS cells treated with DRB in the presence of absence of sgRNA against BLM. For representative images, see Figure S4C. An unpaired t-test was used for statistical analysis; P ≤ 0.01 indicated as **, P ≤ 0.0001 indicated as **** on the graphs.

Deletion of DDX39A did not negatively impact the proliferation of U2OS or the ALT-negative cell line HeLa (Figures S3A-E). DDX39A-depleted U2OS cells displayed higher C-circle levels, as detected by RCA (Figures 3C and 3D) and maintained high levels of telomeric ssDNA (Figures 3E and 3F). The complementation of DDX39A-null cells with DDX39A restored the ssTelo signal to levels similar to that of the parental U2OS cell line (Figure 3F). Notably, the depletion of DDX39A in HeLa cells did not result in an increase in ssTelo signal (Figure 3E). These data confirm the results of our screen and identify the RNA helicase DDX39A as a novel ALT-suppressor.

DDX39A has a closely related paralog, DDX39B, that was not present in our library, and we hypothesized that DDX39B might play a similar role to DDX39A at telomeres in ALT-positive cells. DDX39B was detected in the telomeric chromatin of ALT-positive and negative cells in multiple studies^53, 54^. To test this, we downregulated DDX39B individually or in combination with DDX39A using siRNA (Figures S3F-K). These data showed that the depletion of DDX39B resulted in the accumulation of telomeric ssDNA to a similar extent as DDX39A, raising the possibility that DDX39B might partially compensate for the loss of DDX39A in ALT-positive cells. However, cells co-depleted of DDX39A and DDX39B did not show a further increase in the level of ssDNA (Figures S3I and S3K). Interestingly, contrary to DDX39A, depletion of DDX39B resulted in a severe decrease in cell proliferation regardless of the ALT status of the cells (Figures S3L and S3M). We next examined the dependence of each gene on BLM to determine if the observed signal resulted from canonical ALT and found that in every case, BLM deletion strongly reduced the ssTelo signal (Figure 3G and 3H). Collectively, these experiments showed that DDX39A and its paralog, DDX39B, suppressed ALT-associated accumulation of telomeric ssDNA.

Since most subunits of RNAPII were among the TAILS hits, we next asked whether inhibiting RNAPII would stimulate ALT activity. To test this hypothesis, we used 3 different RNAPII inhibitors with distinct mechanism of action: 5,6-dichloro-1-beta-D-ribofuranosyl-benzimidazole (DRB), flavopiridol (FL; a CDK9 inhibitor) and Triptolide (TP). TP affects transcription initiation, while DRB and FL both affect transcription elongation by inhibiting the phosphorylation of RNAPII either indirectly or directly, respectively. Treatment of U2OS cells with any of these inhibitors resulted in a significant induction of ssDNA at telomeres (Figures 3I, 3J, S4A, and S4B). Similar results were obtained in the ALT-positive cell lines G292 and SAOS-2 using DRB (Figure 3I and 3J).

We next examined whether the increase in ssTelo observed with RNAPII inhibitors was due to canonical ALT. To investigate this, we depleted BLM in U2OS cells using sgRNA for 96 hrs and treated the cells with DRB for 6 hrs (Figures 3K and S4C). BLM depletion significantly reduced the ssTelo signal in DRB-treated U2OS cells, indicating that RNAPII inhibition induces ALT-associated telomeric ssDNA. To determine if ALT-positive cells were more sensitive to inhibition of RNAPII, we treated cells with RNAPII inhibitors at a range of doses and found that they had a marked effect on cell growth regardless of ALT status (Figure S4D and S4E). These data indicate that inhibition of RNAPII elevates the BLM-dependent production of ssDNA and suggest that RNAPII-dependent transcription could be an additional target to explore in targeting ALT cancers.

### ALT-suppressors: Replisome-associated histone chaperones

Loss of ATRX-DAXX-mediated histone H3.3 deposition and elevated localization of DNA replication and repair proteins at telomeres are characteristics of ALT-positive cells^55^. Consistent with the importance of these pathways in maintaining the ALT mechanism, TAILS identified many proteins involved in DNA replication and repair, as well as several genes encoding histones and histone chaperones as key ALT modulators (Figures 1F and 3A). These included H3F3B, one of the two genes encoding the histone variant H3.3; HIRA, a replication-independent chaperone for H3.3; CHAF1A, CHAF1B, and RBBP4 that encode the 3 subunits of the Chromatin Assembly Factor 1 (CAF1) histone chaperone complex; SUPT16H, a subunit of the Facilitates Chromatin Transcription (FACT) complex; ASF1A, a chaperone for H3-H4; and SUPT6H, a histone chaperone implicated in both DNA replication and transcription (Figures S1I and Table 1)^56, 57^. These hits suggested that ALT cells may be particularly sensitive to additional defects in histone deposition or exchange, which occurs most abundantly during DNA replication and transcription. To test this, we depleted CHAF1B, which is involved in replication-dependent nucleosome deposition of H3.1/2-H4, and HIRA, a replication-independent chaperone for H3.3-H4. Depletion of either CHAF1B or HIRA by siRNA led to an upregulation of ssDNA at telomeres in the ALT-positive cell lines U2OS (Figures 4A, 4B, and S5A) and LM216J (Figures S5B and S5C), validating the TAILS results. In non-ALT cells, CHAF1B and HIRA were previously shown to regulate the replication-dependent deposition of H3.1 and the replication-independent deposition of H3.3, respectively^58^. We measured *de novo* deposition of H3.1 or H3.3 in U2OS cells stably expressing SNAP-H3.1 or H3.3 using a quench-chase-pulse strategy. Depletion of CHAF1B specifically impaired the *de novo* deposition of SNAP-H3.1, but not SNAP-H3.3 (Figures 4C and S5D). In contrast, depletion of HIRA strongly impaired H3.3 deposition, but not H3.1, indicating chaperone specificity is maintained in ALT cells (Figure 4C). These data both establish the specificity of the assay for replication-dependent and independent pathways of histone H3 deposition and indicate that telomeric ssDNA is elevated by the perturbation of either pathway in ALT-positive cells.

**Fig 4:**
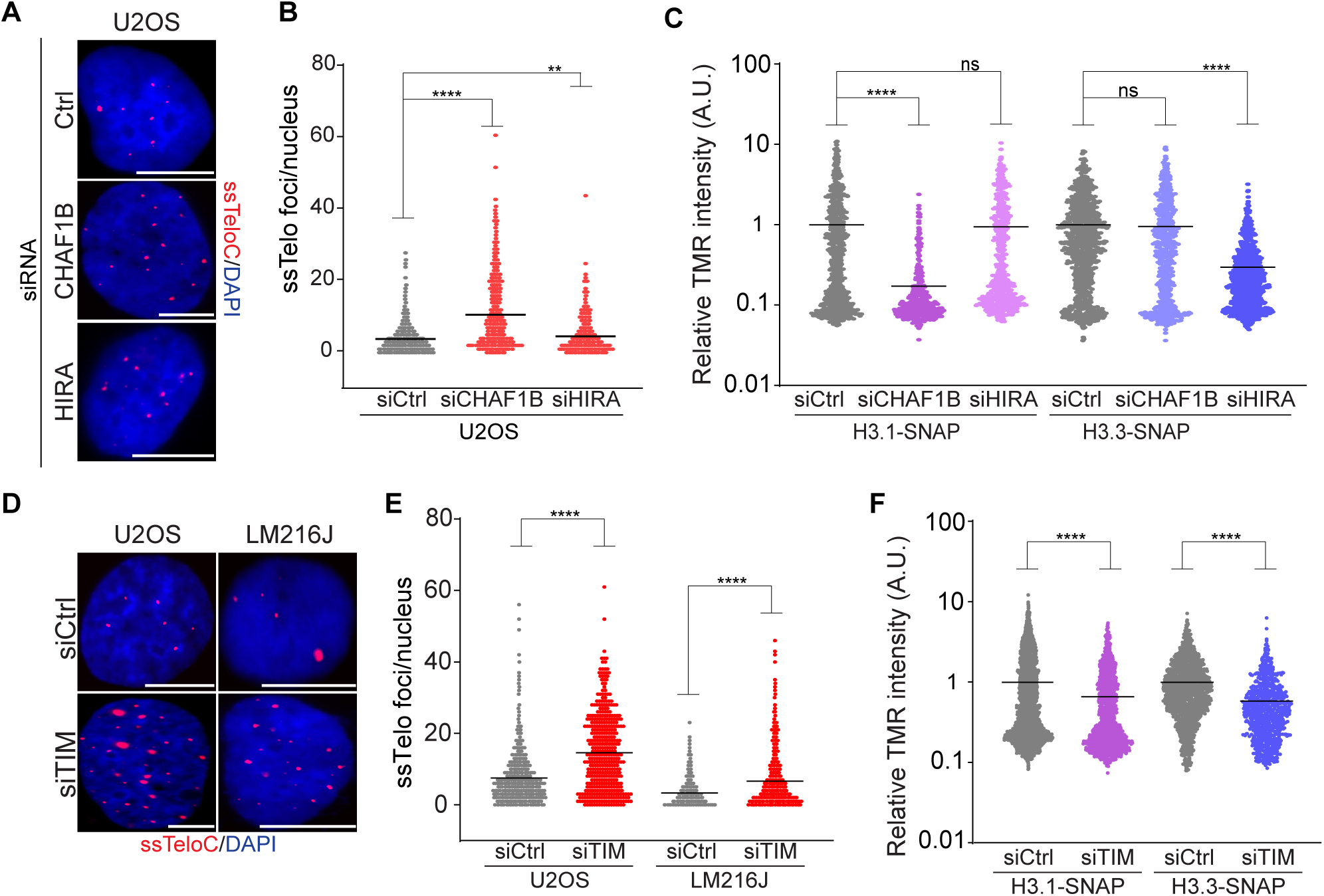
Reduced histone deposition elevates ssDNA at ALT-positive telomeres A-B. Representative images and quantification of ssTelo staining U2OS cells treated with siRNAs against either CHAF1B or HIRA histone chaperones encoding genes. **C** Quantification of *de novo* histone deposition in U2OS cells treated with siRNAs against either CHAF1B, HIRA or a non-targeting control (siCtrl). TMR intensity levels are quantified and averaged, normalized to samples treated with siCtrl, and plotted on a log scale. **D-E** Representative images and quantification of ssTelo staining U2OS cells treated with siRNAs against TIMELESS (siTIM) compared to control cells (siCtrl). **F** Quantification of *de novo* histone deposition in U2OS cells treated with siRNAs against TIMELESS. The scale bar is 10 μm. An unpaired t-test was used for statistical analysis; P > 0.05 indicated as ns, P ≤ 0.01 indicated as **, P ≤ 0.0001 indicated as **** on the graphs.

We next examined the replisome component TIMELESS, which has been implicated in DNA fork protection and, more recently, in the prevention of replication-transcription conflicts^59–61^. siRNA-mediated TIMELESS depletion resulted in increased levels of ssTelo in the ALT-positive cells U2OS and LM216J, validating our TAILS screen result (Figure S5E, 4D and 4E). Structural and functional studies have implicated TIMELESS in the functions of the FACT histone chaperone, and it was proposed to act as a platform to facilitate histone eviction based on structural studies in yeast^62, 63^. We, therefore, asked whether TIMELESS depletion was also associated with defects in histone deposition using the SNAP-tag approach. We found a significant reduction, albeit less severe than perturbation of CHAF1B or HIRA, in the levels of both H3.1 and H3.3 deposition following TIMELESS depletion (Figure 4F and S5F).

The ASF1A and ASF1B histone chaperones are regulated by the Tousled-like kinases 1 and 2 (TLK1 and TLK2). Depletion of TLK1 and TLK2 phenocopies the loss of ASF1A/B in many respects, including impaired H3.1 and H3.3 deposition, as well as the induction of ALT features in both ALT and non-ALT cells^64, 65^. We, therefore, tested the TLK inhibitor E804-20 (TLKi), developed through modifications of indirubin E804, which we previously identified^66, 67^. Following treatment of a panel of ALT-positive and ALT-negative cells with TLKi for 6 hrs, we observed a dramatic elevation in ssTelo only in ALT-positive cell lines (Figures 5A-C). Similar effects were also observed for ATRi (AZ20) (Figures 5A-C), confirming the data obtained by TAILS, which identified ATR as one of the top hits involved in suppressing ssTelo in ALT-positive cells (Figure 3B).

**Fig 5:**
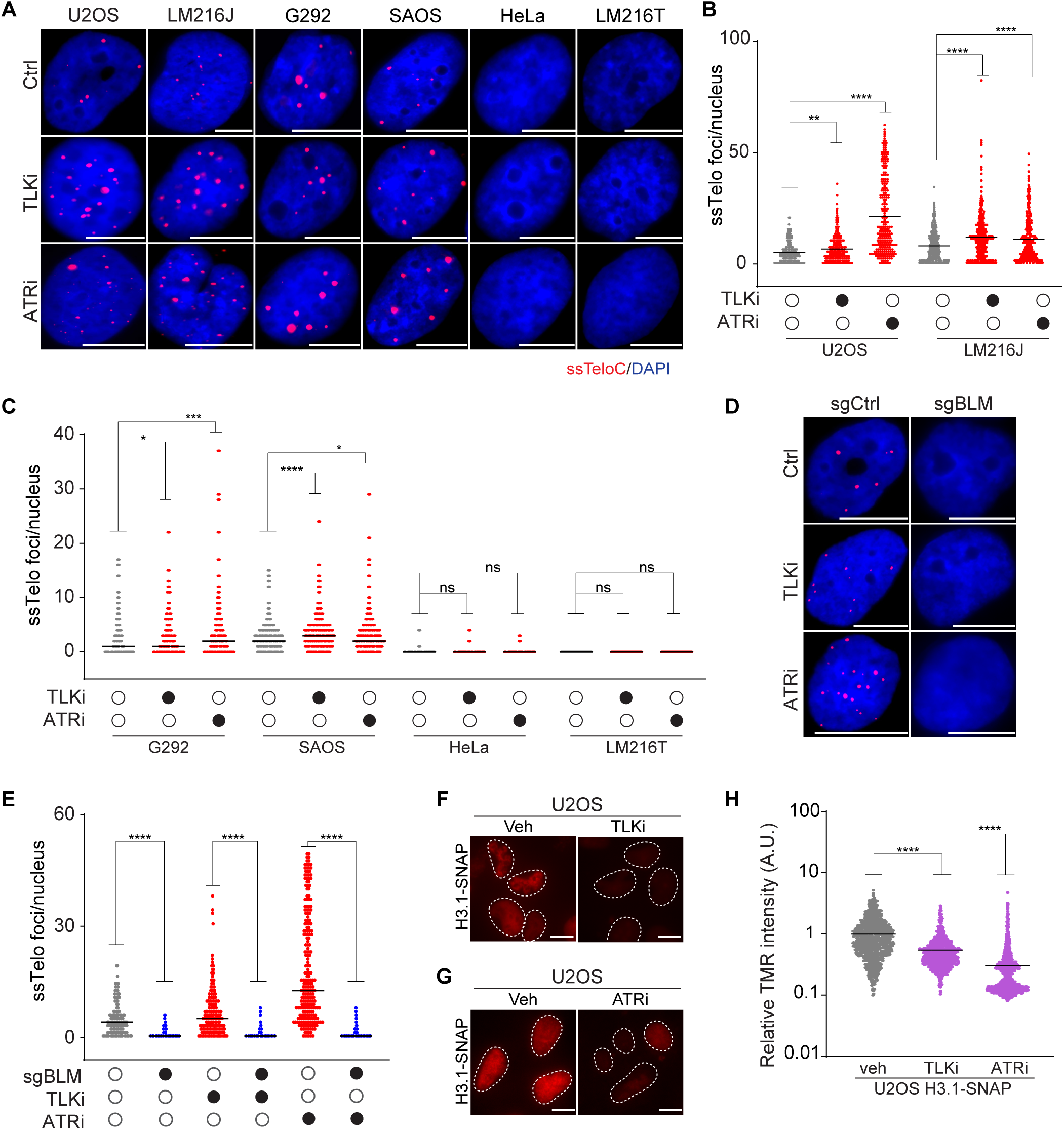
Inhibition of ATR or TLK elevates telomeric ssDNA and reduces histone deposition A-C. Representative images and quantification of ssTelo staining of a panel of ALT-positive (U2OS, LM216J, G292, and SAOS-2) and ALT-negative (HeLa and LM216T) cells treated with TLKi or ATRi for 6 hrs. **D-E** Representative images and quantifications of ssTelo staining of U2OS in the presence or absence of sgRNA against BLM treated with TLKi or ATRi. **F-G** Representative images TMR staining (red) of H3.1-SNAP U2OS cells treated TLKi (F), ATRi (G) or control (veh). **H** Quantification of *de novo* H3.1 histone deposition in U2OS cells treated with TLKi or ATRi. The scale bar is 10 μm. An unpaired t-test was used for statistical analysis; P > 0.05 indicated as ns, P ≤ 0.05 indicated as *, P ≤ 0.001 indicated as ***, P ≤ 0.0001 indicated as **** on the graphs.

We also checked whether inhibition of either TLK1/2 or ATR affected APBs in U2OS cells. We observed no significant effect on APB levels upon these acute treatments (Figures S6A and B). Given that depletion of TLK induces replication stress and is synthetically lethal with checkpoint inhibitors, we wondered whether TLK and ATR inhibition had a synergistic effect on ALT features^68^. To this end, we treated U2OS cells with TLKi and ATRi for 6 hrs and found that co-inhibition of TLK and ATR did not further increase the levels of ssTelo compared to cells treated with only TLKi, suggesting an epistatic relationship between TLK and ATR activity (Figure S6C-S6D). We next tested whether the increase in the ssTelo signal was BLM-dependent by depleting BLM in U2OS cells for 96 hrs and treating the cells with either TLKi or ATRi for 6 hrs. We found that in both cases, depletion of BLM strongly reduced ssTelo signal (Figure 5D and 5E).

We next tested the effects of TLKi and ATRi on histone deposition using the *de novo* histone incorporation assay in U2OS cells. In both cases, increased ssTelo foci correlated with a significant impairment in the deposition of both H3.1 and H3.3 variants (Figure 5F-H and S6E-G). Given that both ATR and TLK1/2 suppress replication stress, we asked whether the elevated ssDNA seen at telomeres was specific to these repetitive regions or was a global effect. Staining with a centromeric probe showed that, while centromeric and telomeric sequences can be readily detected under denaturing conditions, only telomeric repeats displayed ssDNA upon ATR or TLK1/2 inhibition, highlighting the specific effect of ATR and TLK1/2 inhibition on ALT telomeres (Figures S6H and S6I).

Collectively, our data show that ALT features can be stimulated using pharmacological treatments that impact both DNA replication and transcription (RNAPIIi, TLKi and ATRi), two major processes that involve the removal and reestablishment of nucleosomes in chromatin. Depletion/inhibition of TLK1/2, depletion of TIMELESS or depletion/inhibition of ATR caused replication stress, which we demonstrate to be accompanied by the impairment of both replication-dependent and -independent histone deposition pathways ^64, 68–70^. As RNAPII-mediated transcription also involves histone removal and replacement via HIRA, FACT, ASF1A, and SPT6, all of which were identified as hits by TAILS (Supplemental Table1), this further supports the idea that ALT cells may be similarly affected by multiple perturbations that reduce histone content^71^.

## Discussion

Here, we report the development and implementation of a high-throughput imaging screen, termed TAILS (Telomeric ALT *In situ* Localization Screen), to identify cellular factors regulating the ALT pathway. Using this approach with a focused CRISPR-KO sgRNA library, we identified over 80 new factors that influence the generation of ssDNA at telomeres in ALT cells. This approach primarily identified factors that suppress ALT activity, along with a handful of factors that promote ALT activity. This could reflect either a lower resolution in detecting loss of signal or the underlying biology of the ALT process.

In addition to BLM, PML, and members of the SUMOylation pathway, TAILS identified PPP2CA, a protein-phosphatase subunit, and three chromatin remodeling factors, CHD4, EP300, and SGF29, as factors that reduce ALT activity when knocked-out. Although we did not further validate EP300 or PPP2CA here, previous studies have shown that PPP2CA regulates several proteins implicated in the DDR and ALT, including ATM, CHK1/2, and TLK2^64, 72^. PPP2CA has also been implicated in the generation of TERRA via the regulation of hnRNPA1^73^. Recent work also established a role for EP300 as an ALT-promoter using siRNA and small molecule inhibitors consistent with our TAILS results^74^.

Depletion of the NuRD complex component CHD4 or the SAGA/ATAC complex component SGF29 significantly reduced ssTelo signal in multiple ALT cell lines, validating our screen and showing that these factors are indeed required for the ALT process. The results of our screen indicated that deletion of the rest of the core components of the NuRD complex did not significantly alter the amount of ssTelo signal, with the exception of RBBP4, which led to a strong increase in ssTelo signal, likely due to its role in the CAF1 complex (Figures S1I and Supplemental Table 1). Previous work demonstrated that the NuRD complex is recruited to ALT telomeres by ZNF287 and it plays a significant role in the recruitment of HR factors^43^. CHD4 contains tandem PHD domains and tandem chromodomains that are thought to recognize several histone marks associated with transcriptionally inactive chromatin^75^. Future work will be needed to establish the precise role of CHD4 in the ALT process. As multiple inhibitors for histone-binding domains have been developed, CHD4 is also a potentially targetable modulator of ALT.

We validated the SAGA/ATAC complex component SGF29 as another novel promoter of ALT (Figure 2C-D). In contrast to the NuRD complex, the SAGA and ATAC complexes are associated with transcriptionally active chromatin through the recognition of H3K9ac and H3K14ac^41, 76^. SGF29 is well established to be a component of the HAT modules of both the SAGA complex, that contains the HAT KAT2A/GCN5, and ATAC, that contains either KAT2A/GCN5 or KAT2B/PCAF HAT activities^41^. Both KAT2A and KAT2B were present in the screen and their individual loss did not lead to a notable reduction in ssTelo intensity (Supplementary Table 1). Previous work in human embryonic stem cells showed that the loss of the SAGA or ATAC complexes impaired specific sets of genes involved in self-renewal, independently of the HAT activities of either complex^77^. Our data suggests that either the loss of multiple HAT functions, through the perturbation of KAT2A and KAT2B complexes, or a HAT-independent function of SGF29 may be important in ALT. Notably, few components of the ATAC or SAGA complexes have been identified in telomeric chromatin in proteomic approaches^53, 54^. Thus, it remains to be determined whether the effects of SGF29 in ALT are direct, through recognition of telomeric chromatin, or indirect, via effects on gene expression, and whether targeting SGF29 could be effective or selective in ALT-positive cancers.

TAILS identified many new ALT-suppressors, including the putative RNA helicase DDX39A (Fig. 3). In addition, we observed a similar phenotype when we knocked out its closely related paralog, DDX39B. Both helicases have been shown to localize to telomeres and are associated with the THO/TREX complex, which suppresses the accumulation of R-loops^51, 53, 54, 78, 79^. DDX39B depletion was shown to facilitate nascent RNA processing to prevent R-loop formation, and its loss impaired transcriptional elongation, induced hyper-recombination, caused DNA damage, and led to genome-wide R-loop accumulation in non-ALT cells^80, 81^. In ALT-positive cells, the THO/TREX complex was shown to prevent TERRA accumulation and telomere fragility, although a role for DDX39B in this process was not established^82^. Based on these previous studies, we speculate that depleting these helicases could further increase the levels of TERRA R-loops that are present at high levels at ALT telomeres, stimulating the ALT process^83^. Another non-exclusive possibility is that the loss of DDX39A and/or DDX39B impairs the splicing of genes critical for telomere maintenance^84^. Moreover, a recent study found that in mouse embryonic stem cells DDX39, that is a single gene in mice, can be recruited to telomeres in a TRF1-dependent manner to modulate telomere length^85^. Further work is required to uncover the specific direct or indirect roles of DDX39A/B at telomeres in ALT-positive cancer cells.

Our work highlights a novel role for TIMELESS in limiting ALT activity at telomeres (Fig. 4D-E). Previous work showed that the TIMELESS/TIPIN complex suppresses telomere clustering and telomeric MiDAS in ALT cells^86^. Here, we demonstrate that TIMELESS depletion caused accumulation of telomeric ssDNA and reduced histone deposition, suggesting a critical role for this factor in maintaining telomeres in ALT cells. TIMELESS alleviates replication stress and prevents hyper-resection at stalled forks by promoting RAD51 loading^87^. Recent studies identified TIMELESS and TIPIN as crucial for resolving transcription-replication conflicts, functioning in an epistatic fashion with PARP1^61^. While TIMELESS was a clear hit in TAILS analysis, PARP1 and TIPIN did not score as strongly, suggesting a potentially distinct role for TIMELESS in ALT suppression. Additionally, we found that TIMELESS promotes *de novo* histone deposition, an ASF1-dependent process that deposits hypo-methylated histones that are critical for RAD51 loading by the TONSL-MMS22L complex^88, 89^. TIMELESS also promotes the resolution of G4 structures through the recruitment of DDX11, a function that could also be relevant to the ALT process^90^. Therefore, the loss of TIMELESS could lead to elevated ssDNA in ALT-positive cells through multiple mechanisms.

Another major class of factors identified by TAILS as ALT-suppressors were subunits of the RNAPII complex. We validated these findings by demonstrating that chemical inhibition of RNAPII activity resulted in the accumulation of ALT features (Figures 3I, 3J, S4A and S4B). This can be explained by several potential mechanisms. One possibility is that suppression of RNAPII results in frequent transcription-replication conflicts at telomeres, leading to replication stress and R-loops, that cause DNA damage and increased ALT activity. RNAPII-mediated transcription also impacts chromatin structure at telomeres through histone exchange and the recruitment of histone modifying enzymes^91, 92^. Given the identification of multiple RNAPII-associated histone chaperones, including HIRA, SPT6 and FACT, as ALT-suppressors, this suggests a model linking RNAPII with histone chaperone activity at telomeres in ALT cells^71^.

Finally, our data shows that ALT activity can be rapidly modulated pharmacologically using a number of small molecules, including inhibitors targeting RNAPII, TLK, ATR, and SUMOylation inhibitor. To date a handful of small molecules have been shown to modify the ALT process, including inhibitors of ATR, PARP, Topoisomerases, HDACs, Bromodomain proteins, the ABL1-JNK-JUN signaling pathway, and SUMOylation^23, 30, 49, 93, 94^. Using predictions from our TAILS screen, we identified more than eight additional small molecules that modulate ALT features, including compounds that either increased or decreased the number and intensity of ssTelo signals at telomeres. Previous work reported a reduction in ALT features following extended ATR inhibition (24-48 hrs), including lower levels of C-circles, APBs and T-SCE^23^. In contrast, here we found that acute treatment (6 hrs) with ATR inhibitors markedly elevated the levels of ssTelo signal in multiple ALT cells lines. These differences in outcome may reflect the timing or off target effects of the different small molecules used and further analysis will be required to understand the underlying effects of ATR inhibition on ALT.

ATR inhibition was proposed to be selectively toxic for ALT-positive cells, although these results remain controversial, as ATR inhibitors did not clearly show selective toxicity in ALT-positive cells in subsequent studies^23, 95, 96^. While TLK depletion and inhibition modify many features of the ALT process, we did not observe ALT-selectivity in the few cell lines we analyzed. In data from the DepMap project^97^, the loss of TLK2 reduced the fitness of the majority of cancer cell lines tested, similar to ATR, and TLK2 deletion in mouse models of AML markedly impaired cancer growth, indicating that TLK inhibition will likely have broader anti-cancer effects that are not limited to ALT-positive cells^98^. Although future work will be needed to identify additional small molecules, establishment of the TAILS approach provides a powerful method for the future identification of potentially selective compounds to treat ALT cancers.

### Limitations of the study

This study provides valuable novel insights into factors that modulate ALT activity, yet it has some limitations. We performed our screen in a single ALT-positive cells line (U2OS) and validated our results for a selected pool of factors in various independent cell lines. It remains possible that some of the genes identified may have cell line specific effects on ALT. Additionally, an intrinsic limitation of high throughput screens is the potential for false positive and negative hits. Further work will be needed to address these aspects and strengthen the generalizability of the findings. Finally, while we used multiple assays to validate a selection of hits, the ssTelo readout may not always correlate with all aspects of the ALT process and further analysis of individual hits will be needed to establish their precise roles in ALT.

## Acknowledgements

We are grateful to Tom Misteli for providing reagents and Laurent Ozbun for his technical help with the screen. We thank members of the Lazzerini Denchi and the Stracker labs, and Sam John for critical feedback on this work, and Roderick O’Sullivan (UPitt) for discussion, reagents and sharing unpublished data. Research in the ELD and THS labs and at HiTIF are funded by the NIH Intramural Research Program of the National Institutes of Health (NIH), National Cancer Institute (NCI) and Center for Cancer Research, projects 1-ZIA-BC011815-03, 1-ZIA-BC 012010-04 and 1-ZIC-BC011567-09, respectively. KMM is supported by NIH NCI (RO1 CA198279 and CA250905) and Cancer Prevention and Research Institute of Texas (RP220330).

## Author contributions

B.A. and E.L.D. designed the study; B.A. performed the TAILS screen; G.P. and B.A. performed the bioinformatic analyses of the TAILS screen; B.A., S.K., S-C.W., G.M.T, S.S, C.J, Y.C performed the experiments; E.L.D, T.H.S. and K.M.M. supervised the project; B.A, T.H.S and E.L.D wrote the manuscript.

## Materials and Methods

### Cell culture

Cell lines were grown in DMEM medium supplemented with 10% fetal bovine serum (FBS) and 0.5 mg/ml penicillin, streptomycin and L-glutamine (Gibco) and cultured at 37°C and 5% CO_2_. G292 cells were grown in McCoy medium supplement with 15% FBS and 0.5 mg/ml penicillin, streptomycin and L-glutamine (Gibco). Cell lines were routinely tested for mycoplasma contamination. LM216J and LM216T cells were a kind gift from Roderick J. O’Sullivan (University of Pittsburgh).

### Generation of DDX39 knockouts

Knockout clones were generated via CRISPR/Cas9 gene targeting as previously described^19^. The following sgRNAs were expressed using a pCDNA-H1-sgRNA vector (Addgene): DDX39A guide 1 (5’-CTCAAAGCCACAGTCCACGA-3’) and guide 2 (5’-TCTTTTGGATTACGATGAAG-3’).

### Cell proliferation analysis

To measure cell proliferation, cells were plated in 6-, 12- or 96-well plates and analyzed using an Incucyte S3 live-cell analysis system (Essen Biosciences) based on confluency. Imaging was performed at 4 hours intervals using a 10x objective. Each condition was normalized to its respective starting confluency at the first timepoint.

### Lentiviral-mediated gene expression

Lentiviral particles were generated by transfection into HEK293T followed by transduction as previously described^19^. The following constructs were used: pLentiCas9-Blast (Addgene) and pBABE-Blast-SNAP-H3.1 or H3.3 vectors (a kind gift from L. Jansen^99^). For SGF29 shRNA-mediated gene silencing the following shRNA were cloned into the pRSITEP-U6Tet-sh-EF1-TetRep-2A-Puro plasmid (Cellecta):

shSGF29-1:CGCGGAAATCAAGTCTCTGTT,

shSGF29-2: CCTGTTTGAAGACACCTCCTA,

shSGF29-3: CCTCAATGTGGCTCAGAGATA.

### Plasmid, siRNA and sgRNA transfections

Wild type and D450A TRF1-FokI expressing mCherry-ER-DD-TRF1-FokI plasmids were a kind gift from R.J. O’Sullivan (University of Pittsburgh)^48^. CHD4 cDNA was cloned into a pcDNA6.2 N-EmGFP-DEST vector. Cells were sequentially transfected with GFP-CHD4 and mCherry-TRF1-FokI plasmids 48 and 24 hours prior to fixation, respectively.

#### siRNA transfections

Transfection of siRNA (final concentration of 50 nM) was carried out either with DharmaFECT1 (Dharmacon) or Lipofectamine RNAiMAX (ThermoFisher Scientific). Cells were harvested 72 hours post transfection.

#### sgRNA transfection

Transfection of sgRNA (final concentration of 2 nM) was carried out with Lipofectamine RNAiMAX (ThermoFisher Scientific) and cells were harvested 96 hours later. Commercially available synthetic sgRNA oligos purchased from Synthego as sgRNA Kits V2 was used to knock out BLM and FANCM genes. Each gene is targeted by a pool of three staggered sgRNA oligos that, when complexed with spCas9, cut in genomic regions 50–100 bp apart. Following sgRNAs were used in this study: BLM (AGAUUUCUUGCAGACUCCGA,UAAAAAUGCUCCAGCAGGAC,UUCACUGAAGGAAAAGUC UU), FANCM (GCUCUGAGGUCGCUCAGUUC,CGAUGAUGUGUUGCUUGUCG, UGGCGGGUUCUGCACCUCCG) and Control non-targeting (GUAACGCGAACUACGCGGGU).

### Small molecule inhibitors

The following inhibitors were used in this study: TLK1/2 inhibitor E804-20^66^ (final concentration of 5 µM in DMSO); ATR inhibitor (final concentration of 10 µM in DMSO); SUMO inhibitor ML-792 (final concentration of 1 µM in DMSO); DRB (final concentration of 200 µM in H_2_O); Triptolide (final concentration of 1 µM in DMSO); Flavipiridol (final concentration of 5 µM in DMSO). Unless otherwise stated, treatments were carried out for 6 hrs before cell harvesting.

### qRT-PCR

Total RNA was extracted from cells using the RNeasy Plus Mini Kit (Qiagen), and cDNA was made using SuperScript II Reverse Transcriptase (ThermoFisher). qPCR was performed using Power SYBR Green PCR Master (Fisher Scientific). Samples were run on a BioRad CFX Opus 96 and analyzed on BioRad CFX Maestro. See key resources table for the sequences of oligonucleotides used in these experiments.

### Immunofluorescence and fluorescence in situ hybridization

IF, FISH, ssTelo staining were carried out as described previously^19^.

### Imaging, image analysis and statistical analysis

Single-plane images were acquired using a Zeiss Axio Imager M2 with an Axiocam 702 camera and a Zeiss Axio observer with an Axiocam 712 camera using ZEN 2.6 (blue edition) software, and an Olympus Fluoview 1000 using FV10-ASW3.1 software. For each experiment and condition, a minimum of 250 cells were imaged. For APB analysis, colocalization was defined as two foci overlapping by 50% or more. For *de novo* histone deposition assay, a Lionheart FX (Agilent BioTek) with Gen5 software was used for analysis. GraphPad Prism v10 was used for statistical analysis and p-values were calculated using the unpaired parametric t-test, unless otherwise stated, the sample size was not predetermined.

### Rolling circle assay (RCA) reaction

RCAs were carried out as described previously^19^ with the following modifications: gDNA was isolated using DNeasy Blood and Tissue gDNA extraction kit (Qiagen), DNA concentrations were determined using the Qubit dsDNA HS assay kit (ThermoFisher) and 30 ng of isolated genomic DNA was used as a template for rolling circle amplification, the product was dot-blotted, UV-cross-linked, and hybridized with an IRDye800 conjugated TelC (5’-5IRD800-CCCTAACCCTAACCCTAA-3’) probe (IDT) with rotation overnight at 37°C. The membrane was scanned and analyzed using the Li-COR Odyssey CLx Imaging System.

### *De novo* histone deposition assay

*De novo* histone deposition assay by quench-chase-pulse was carried out as described previously^58^. U2OS cells expressing SNAP-H3.1 or SNAP-H3.3 were incubated with 5 µM SNAP-Cell Block (NEB) for 30 min at 37°C for the quench step. Cells were washed with PBS twice and incubated in fresh media for 30 min. Following this, cells were again incubated in fresh media for the chase step of 6-7 hours. Lastly, cells were incubated with 1 µM TMR (tetramethylrhodamine) star (NEB) for 30 min at 37°C for the pulse step. Finally, cells were washed and incubated in fresh media for 30 min 37°C, followed by two more PBS washes. The cells were pre-extracted with 0.2% Triton X-100 (Sigma) for 5 min on ice, followed by fixation with 4% paraformaldehyde for 10 min at room temperature. Cells were stained with DAPI. For the experiments with ATRi and TLKi inhibitors, the treatments were carried out during the chase step.

### TAILS

The arrayed sgRNA library (Synthego) used in this study targets 1064 human genes involved in DNA transactions (see Supplementary Table 1 for details). Each gene in the library is targeted by a pool of 3 sgRNA oligos in a single well. As controls, we included non-targeting sgRNAs (Negative control) as well as sgRNA pools against PLK1 (Transfection efficiency control) and BLM (ssTelo control). The sequences of the sgRNAs are provided in Supplementary Table 2. sgRNAs (0.08 pmoles/well) were spotted in an empty 384-well imaging plate (CellVis, P384-1.5H-N) using an ECHO525 acoustic liquid handler (BeckmanCoulter). Plates were air dried, sealed, and stored at −20°C until the day of the reverse transfection. Reverse sgRNA transfection, ssTelo staining in 384-well plate, and image acquisition and analysis were carried out as described previously^38^.

### CRISPR-KO Screen Statistical Analysis

Statistical analysis of the screen was performed using R(Version 4.3.2)^100^ and the cellHTS2 package (2.64.0)^101^. Raw values for each well were first normalized on a per plate basis using the B-score algorithm, which uses the median of the sample population, row and column offsets to normalize positional effects, and the median absolute deviation (MAD) of the samples. Z-score was calculated using the robust Z-score method, which uses the median and MAD of B-score distributions for the sample population. Z-scores for each of the 2 replicates were averaged. RStudio was used for data visualization and statistical analysis.

## Key resources table

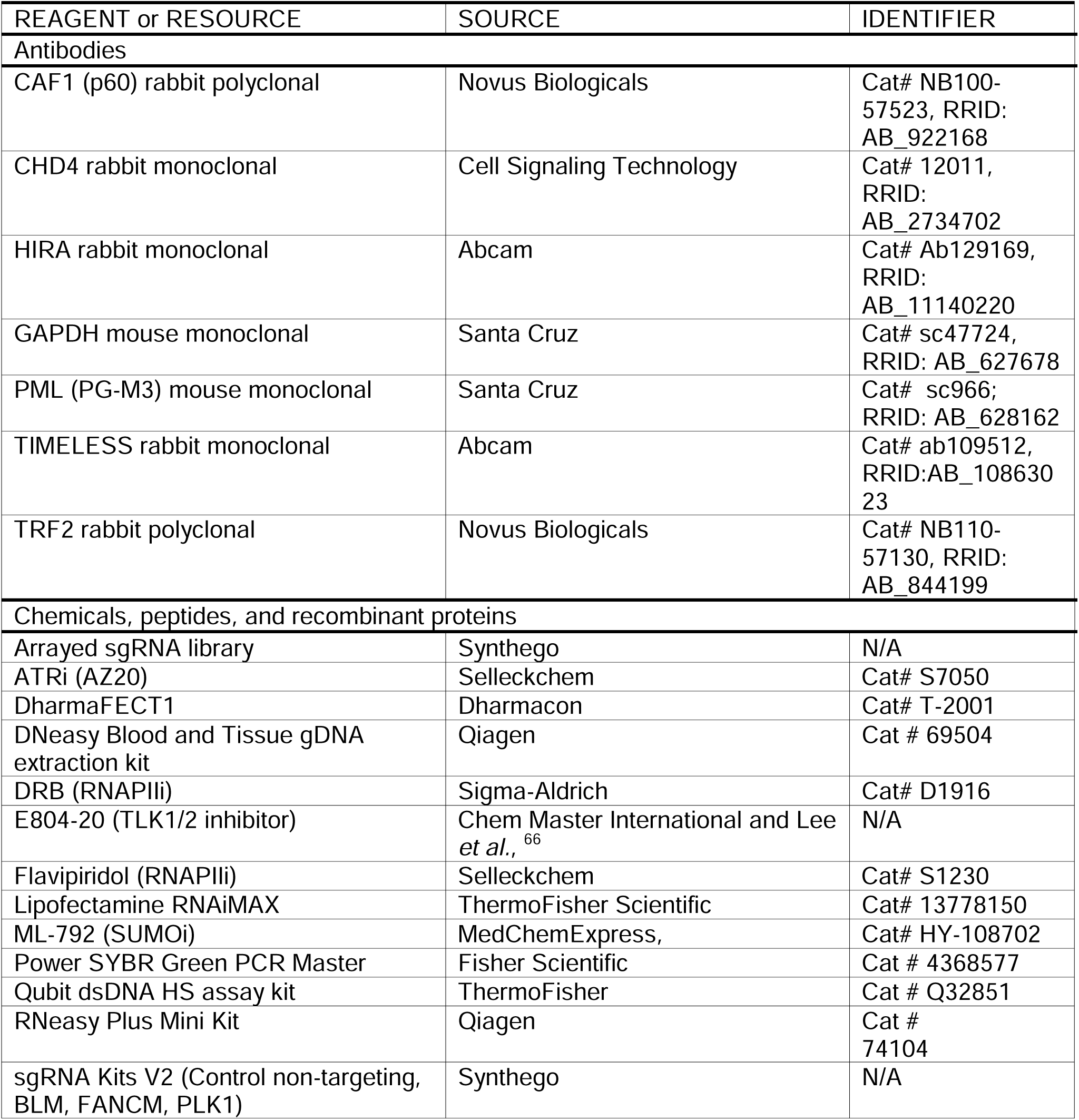
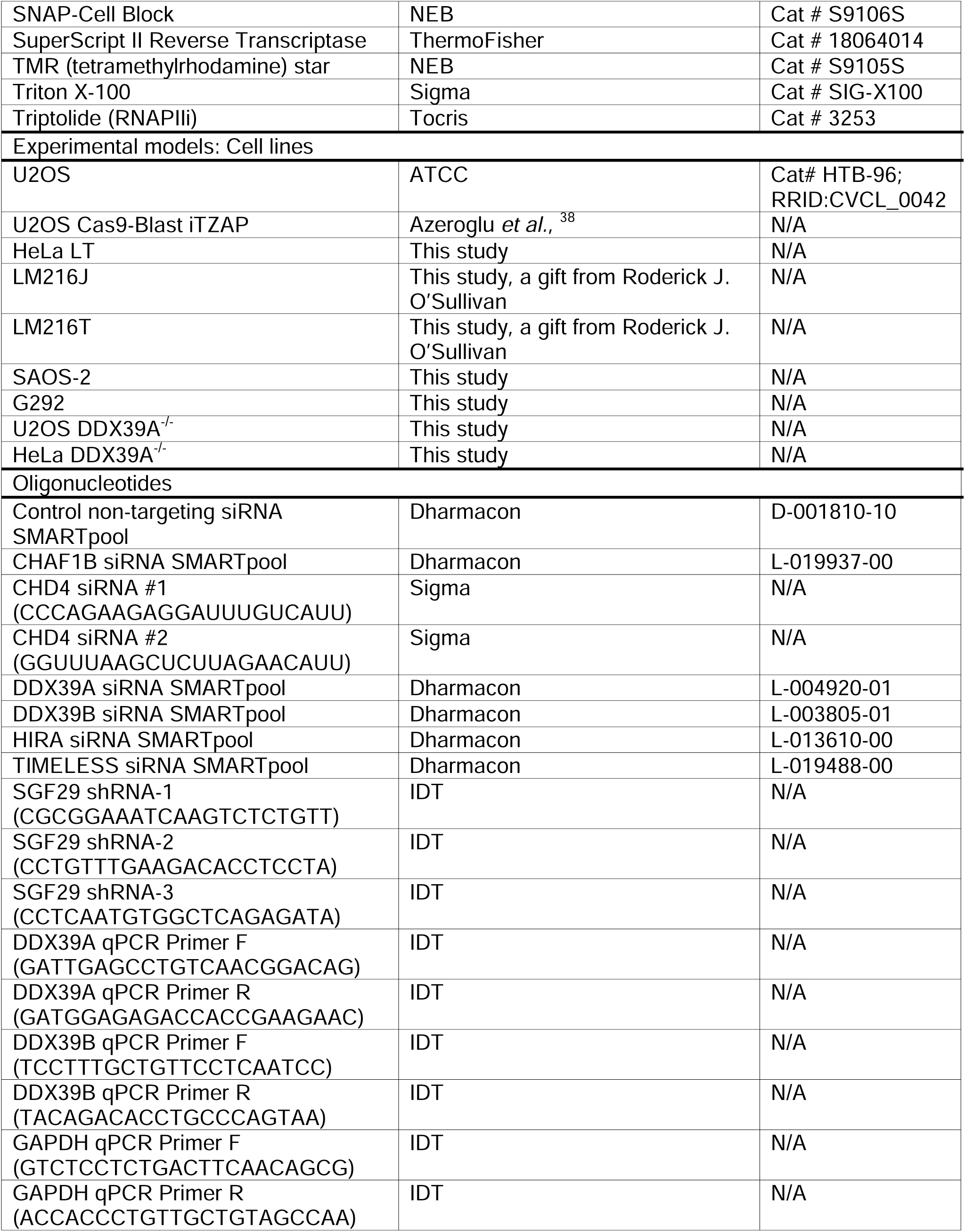
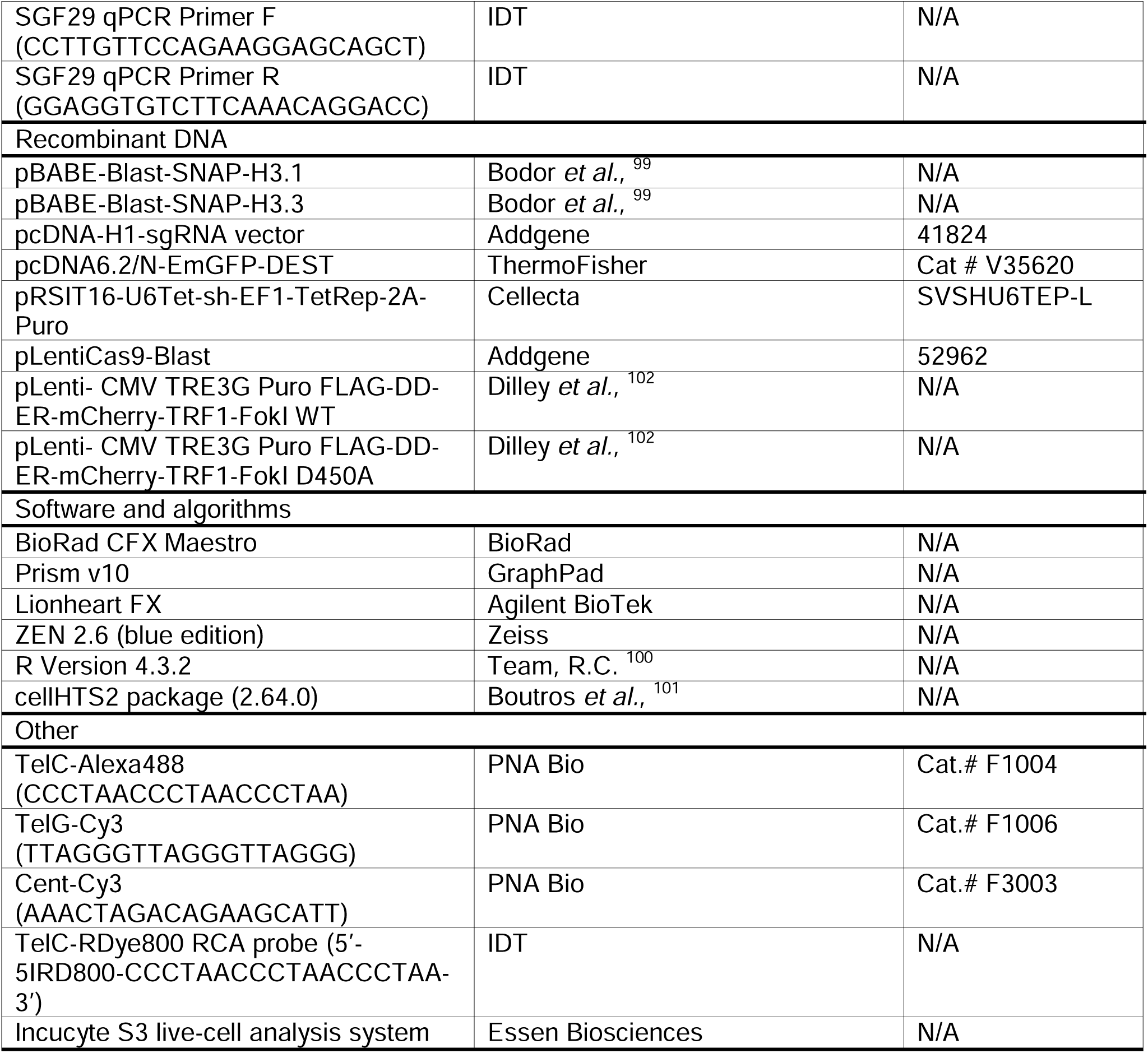

---

**Figure S1.**
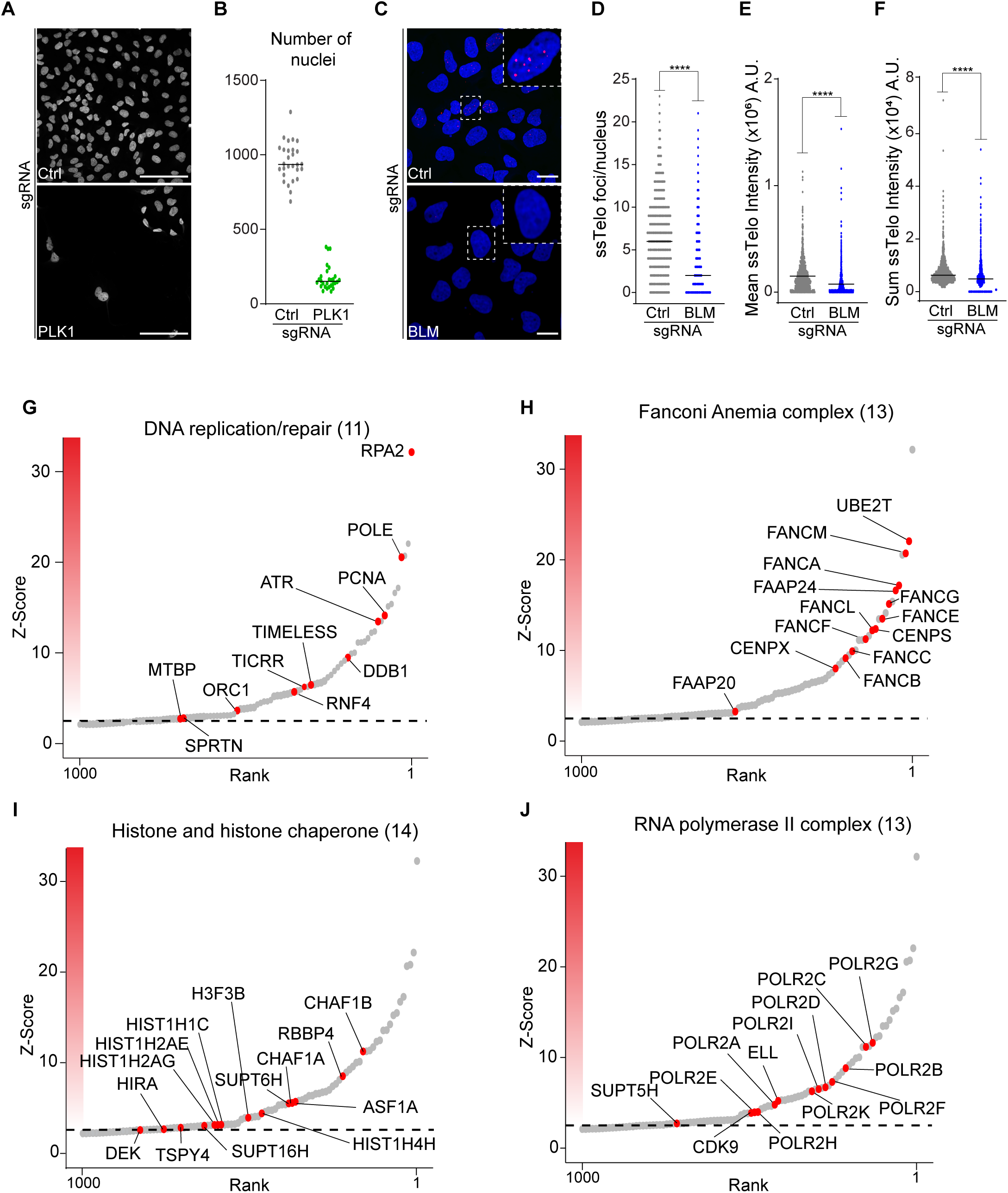

**Figure S2.**
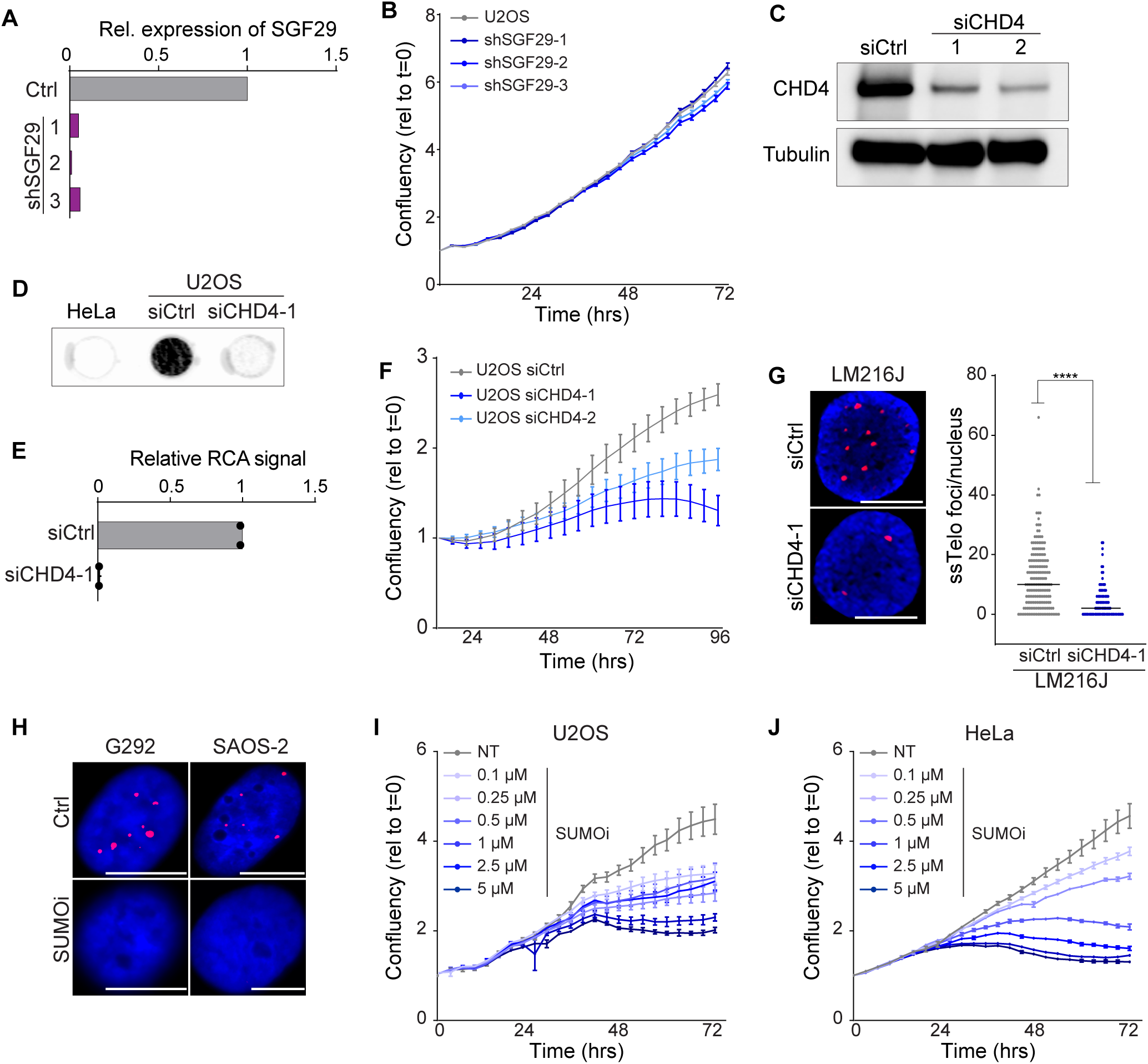

**Figure S3.**
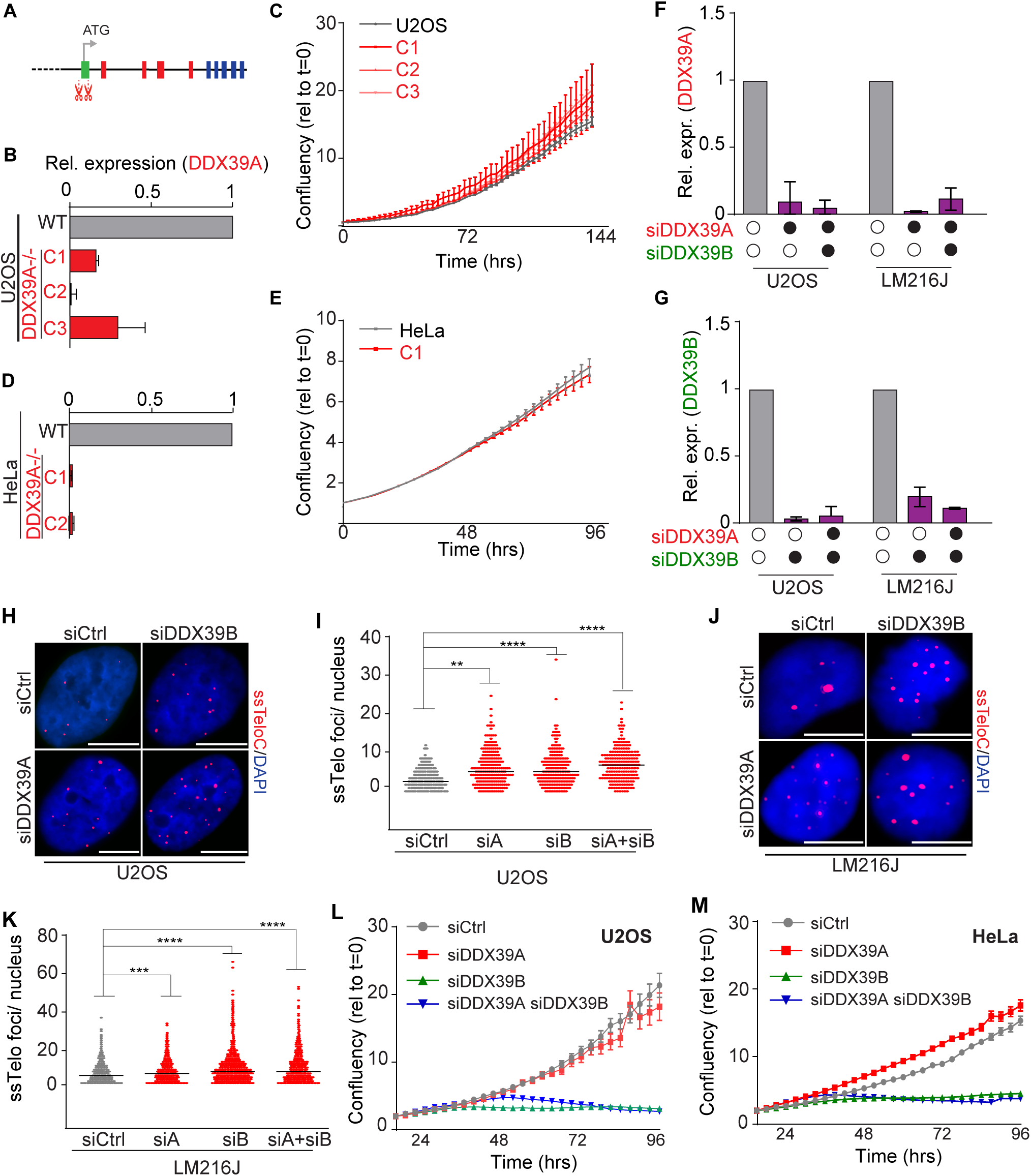

**Figure S4.**
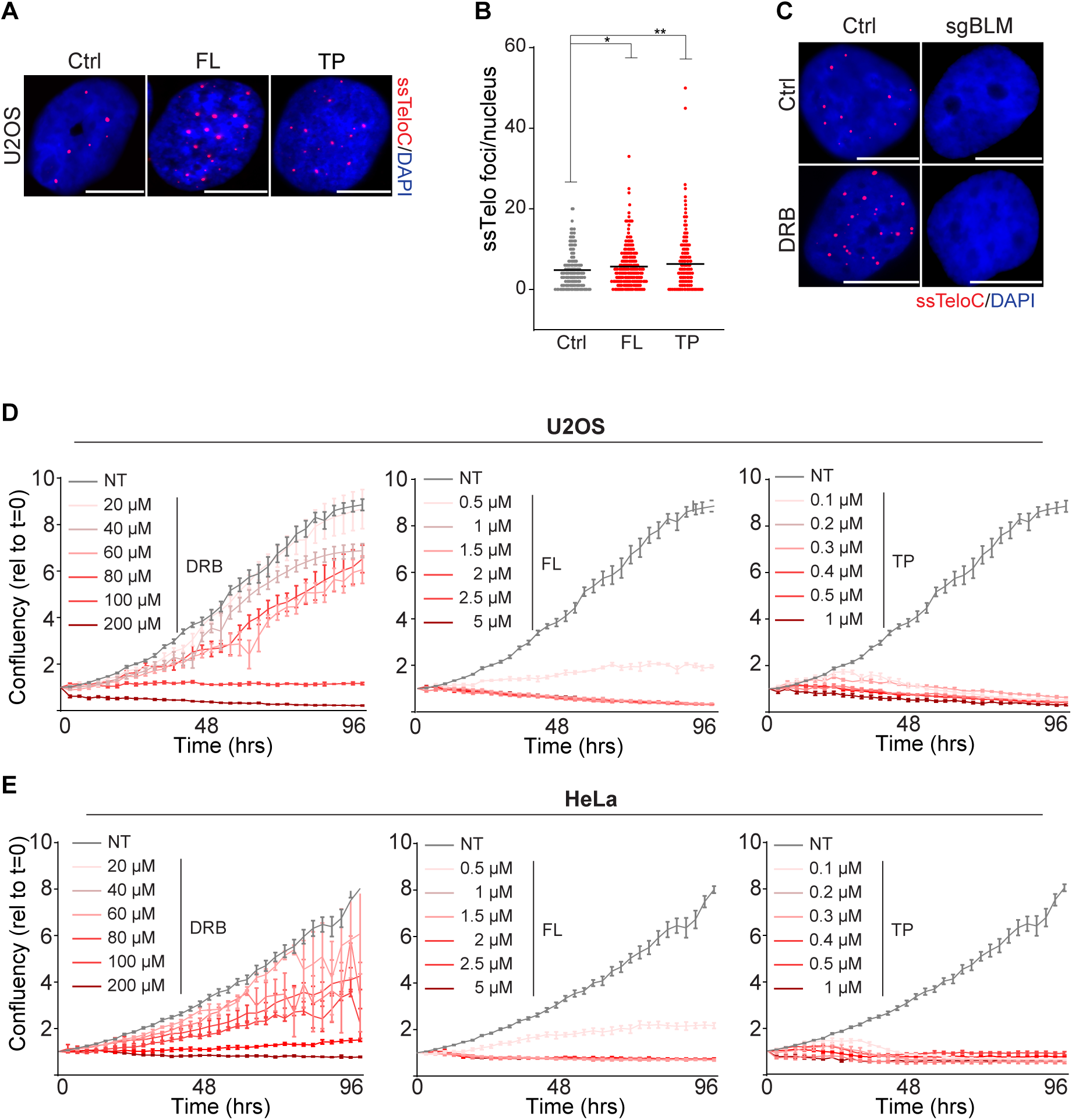

**Figure S5.**
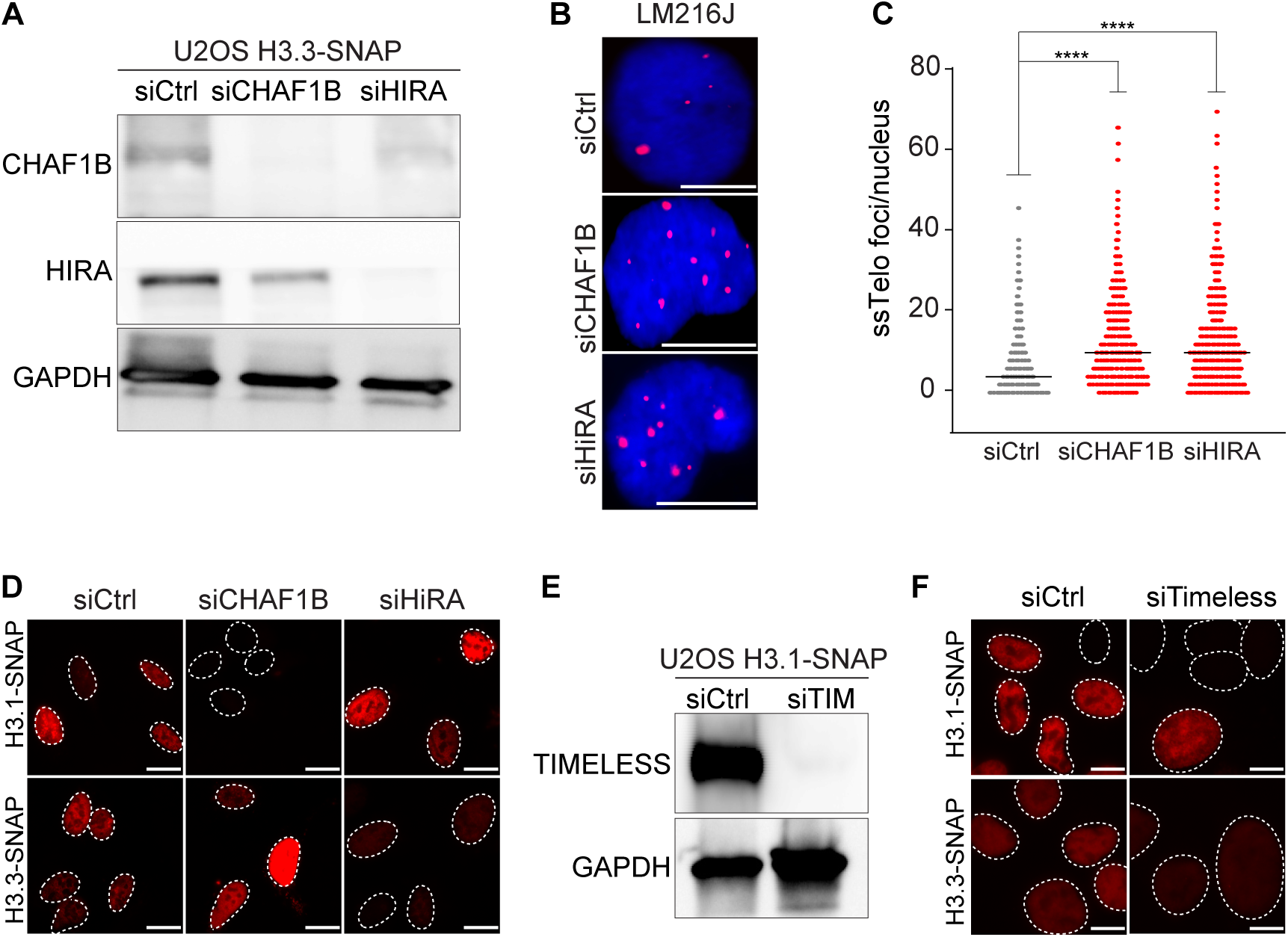

**Figure S6.**
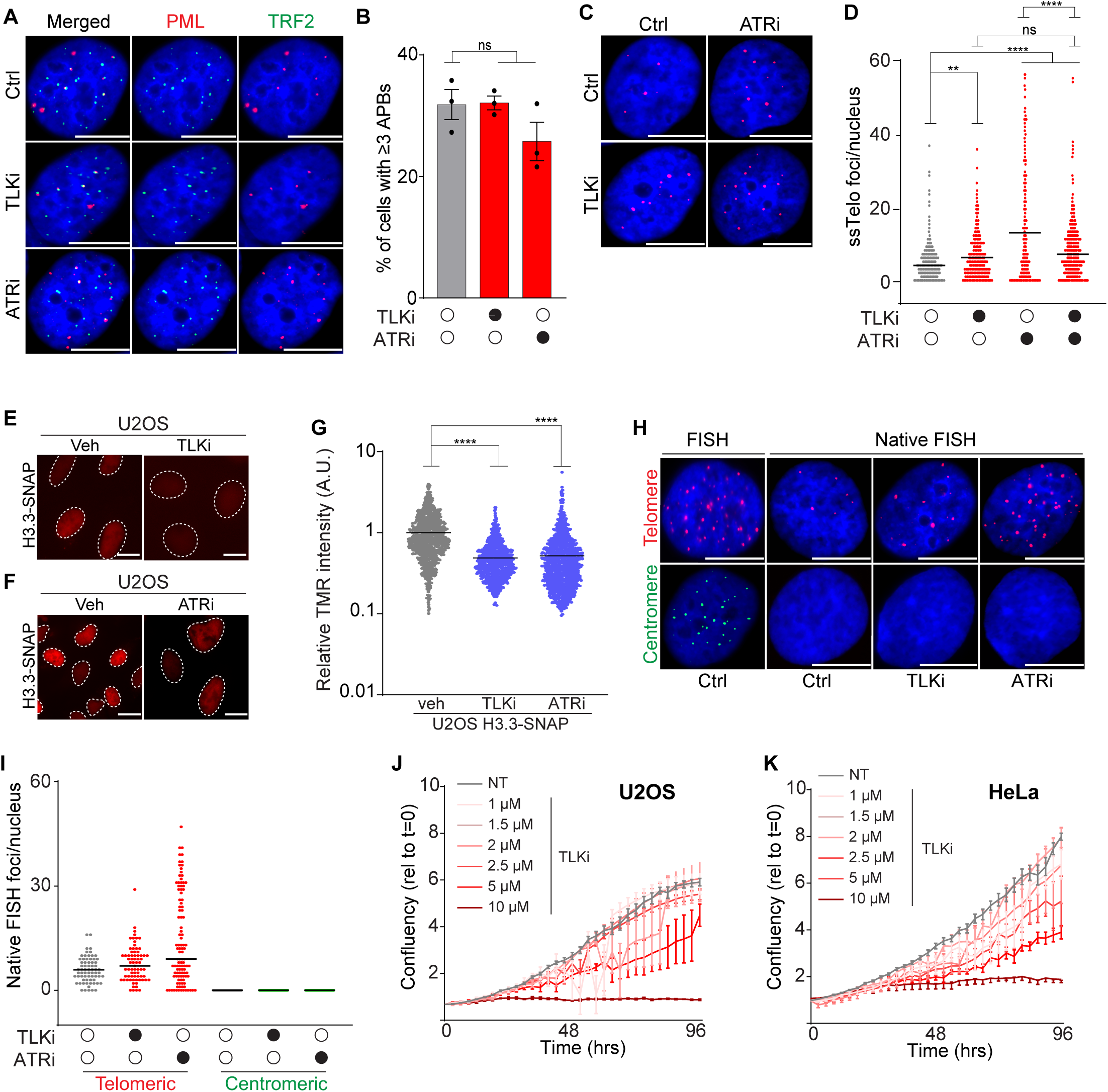

