## Supplementary text for "Identification of Novel Modulators of the ALT Pathway Through a Native FISH-Based Optical Screen"

### Legends to supplementary figures

#### Figure S1 Functional classification of top ALT-inhibitors. Related to Figure 1.

**A** DAPI staining of U2OS cells treated with sgRNA against non-targeting control (Ctrl) or PLK1, 4 days post transfection. The scale bar is 100 μm. **B** Quantification of the data shown in panel A suggest that efficiency to sgRNA-mediated deletion of the essential gene PLK1 is approximately 80%. **C** ssTelo of U2OS cells transfected with a sgRNA against non-targeting control (Ctrl) or BLM. The scale bar is 20 μm. **D-E** Quantification of the data shown in C showing number of ssTelo foci per nuclei (D), mean intensity of ssTelo signal (E) and sum intensity of ssTelo signal (F). **G-J** Scatter plot displaying the average Z-score (y-axis) and gene ranking (x-axis) following TAILS. Gray dots represent genes in the library (n=1064)**,** red dots represent the genes in the library encoding genes involved DNA replication and repair pathway (G), Fanconi Anemia complex (H), histones and histone chaperones (I) and RNA polymerase II complex (H). An unpaired t-test was used for statistical analysis; P ≤ 0.0001 indicated as **** on the graphs.

#### Figure S2 Effect of CHD4 and SGF29-depletion of cell viability. Related to Figure 2.

**A** Expression of SGF29 in U2OS cells expressing 3 different shRNA against SGF29 (shSGF29-1, shSGF29-2, and shSGF29-3) was normalized to U2OS cells (Ctrl) cells. **B** Growth curve of U2OS cells expressing individual shRNA against SGF29 and control. **C** Western blot showing the protein levels for CHD4 and, as a loading control Tubulin in U2OS cells treated with 2 individual siRNA against CHD4 and non-targeting control (siCtrl) 3 days after transfection. **D-E** Rolling circle assay (RCA) analysis of genomic DNA isolated from indicated siRNA-transfected U2OS and HeLa cells. **F** Growth curves of U2OS cells treated with 2 different siRNA against CHD4 or non-targeting control (siCtrl). Growth rate is calculated as confluence relative to confluence at T0. **G** ssTelo staining and relative quantification of LM216J cells transfected with siRNA against CHD4 (siCHD4-1) or a non-targeting control (siCtrl). The scale bar is 10 μm. **H** Representative images of ssTelo staining of G292 and SAOS-2 cells treated with SUMO inhibitor ML-792 (SUMOi). The scale bar is 10 μm. **I-J** Growth curves of U2OS and HeLa across a 6-point dose response for SUMOi tracked every 4 hrs for 72 hrs across a 6-point dose response. X-axis is elapsed time in hours, y-axis is the growth rate as confluence normalized to T0. The null dose is colored grey. An unpaired t-test was used for statistical analysis; P ≤ 0.01 indicated as **, P ≤ 0.0001 indicated as **** on the graphs.

**Figure S3 Characterization of DDX39A/B depletion on cell viability. Related to Figure 3.**

**A** Schematic of the DDX39A genomic locus showing region encoding the DEAD-box RNA helicase domain (green), ATP binding domain (red), and helicase domain (blue). Scissors represent the CRISPR/Cas9 cut sites used to generate knockout clones. **B** Relative expression level of DDX39A in three independent DDX39A$^{-/-}$ U2OS clones compared to DDX39A-proficient U2OS cells (WT). **C** Growth curves of DDX39A$^{-/-}$ clones (C1, C2, and C3) and parental cell line (U2OS). X-axis is elapsed time in hours, y-axis is the growth rate as confluence normalized to T0. **D** Relative expression level of DDX39A in two independent DDX39A$^{-/-}$ HeLa clones compared to DDX39A-proficient HeLa cells (WT). **E** Growth curves of DDX39A$^{-/-}$ clone (C1) and DDX39A-proficient HeLa cells. X-axis is elapsed time in hours, y-axis is the growth rate as confluence normalized to T0. **F-G** Relative expression level of DDX39A and DDX39B in U2OS and LM216J cells treated with indicated siRNAs. **H-I** ssTelo staining and relative quantification of U2OS cells treated with siRNA against DDX39A, DDX39B or a non-targeting control (siCtrl). The scale bar is 10 μm. **J-K** ssTelo staining and relative quantification of LM216J treated with siRNA against DDX39A, DDX39B or a non-targeting control (siCtrl). The scale bar is 10 μm. **L-M** Growth curves of U2OS or HeLa treated with the indicated siRNA. Y-axis is the growth rate as confluence normalized to T0.

**Figure S4 Characterization of RNAPIIi on cell viability. Related to Figure 3.**

**A-B** ssTelo staining and quantification of U2OS cells treated with transcription inhibitors flavopiridol (FL) and triptolide (TP). The scale bar is 10 µm. **C** ssTelo staining of U2OS cells transfected with sgRNAs against BLM and treated with DRB. Related to Figure 3K. The scale bar is 10 µm. **D-E** Growth curves of U2OS or HeLa treated with the indicated doses of the transcription inhibitor DBR, FL and TP. Y-axis is the growth rate as confluence normalized to T0. An unpaired t-test was used for statistical analysis; $P \leq 0.1$ indicated as *, $P \leq 0.001$ indicated as *** on the graphs.

**Figure S5 Defects in histone deposition. Related to Figure 4.**

**A** Western blot on protein lysates of U2OS cells treated with siRNAs against CHAF1B, HIRA or a non-targeting control (siCtrl) harvested 3 days after transfection. GAPDH was used as a loading control. **B** ssTelo staining of LM216J cells treated with siRNAs against CHAF1B or HIRA a non-targeting control (siCtrl). **C** Quantification of the data shown in B. **D** Representative images TMR (tetramethylrhodamine) straining (red) of H3.1-SNAP and H3.3-SNAP U2OS cells treated with siRNAs against CHAF1B, HIRA or a non-targeting control (siCtrl). The white dashed lines represent the nuclear outline. **E** Relative protein levels of TIMELESS in U2OS cells treated with siRNAs against TIMELESS (siTIM) compared to control (siCtrl) 3 days after transfection by Western blot analysis. **F** Representative images TMR straining (red) of H3.1-SNAP and H3.3-SNAP U2OS cells treated with siRNAs against TIMELESS (siTIM) or control (siCtrl). The white dashed lines represent the outlines of the cells. The scale bar is 10 µm. An unpaired t-test was used for statistical analysis; $P \leq 0.0001$ indicated as **** on the graphs.

**Figure S6 Effect of ATRi and TLKi on histone deposition. Related to Figure 5.**

**A** U2OS cells treated with ATRi or TLKi were stained for PML (red) and TRF2 (green). **B** Quantification of data shown in A. Graphs indicate the percentage of cells that have at least 3 PML colocalization to TRF2 (APBs) per nucleus. The colocalization was defined as two foci overlapping by 50% or more. **C-D** ssTelo staining and quantification of U2OS cells treated with TLKi and ATRi either individually or together. One-way ANOVA was used for statistical analysis. **E-G** Representative images TMR straining (red) and quantification of H3.3-SNAP U2OS cells treated TLKi, ATRi or control (veh). **H-I** Representative images and quantification of ALT-positive U2OS cells FISH stained under denaturing and native conditions to label telomeric DNA (Telomere, red) or centromeric DNA (Centromere, green) treated with TLKi (F), ATRi (G) or control (Ctrl). **J-K** Growth curves of U2OS or HeLa tracked every 4 hrs for 96 hrs across a 6-point dose response for TLKi (E804-20). X-axis is elapsed time in hours, y-axis is the growth rate as confluence normalized to T0. The null dose is colored grey. The scale bar is 10 μm. An unpaired t-test was used for statistical analysis; $P > 0.05$ indicated as ns, $P \leq 0.01$ indicated as **, $P \leq 0.0001$ indicated as **** on the graphs.

### Legends for Supplementary Tables

#### Table 1 Library composition and TAILS results, Related to Figure 1.

Rank (column A) and Z-Score (column B) of the genes (column C) were calculated based on two independent biological replicates. For each replicate, individual wells were scored for ssTelo integrated intensity (columns D and E), number of ssTelo foci number (columns H and I) and number of nuclei (columns L and M). The mean (columns F, J, N) and deviation of the two experiments (columns G, K, O) were calculated for each measurement. Genes were classified manually based on their reported function in the following categories (column P): (i) DNA replication and repair, (ii) Cell cycle/mitosis, (iii) Transcription/splicing, (iv) Histone/histone chaperone and (v) Chromatin remodified. Further subclassification (column Q) was carried out based on biological pathways (in total 33) using UniProt and PubMed searches. Genes were further classified based on their previous identification as telomeric proteins in ALT and/or non-ALT cells using the following datasets: proteins from (i) TRF1-BioID pull-down comparison between U2OS and HeLa cells, classified as ALT (column R) using data from [51], (ii) QTIP-iPOND as TelChr (column S) for associated with replicating telomeric chromatin and as TelFra (column T) for causing telomere fragility, using data from [54], (iii) RNA oligonucleotide pulldown as Terra (column U) for TERRA-interacting proteins using data from [103], (iv) telomere PICh of ALT-negative HeLa as PIChH (column V) and of ALT-positive VA13 as PIChV (column W) using data from [53], (v) telomere PICh of MEFs as PIChM (column X) using data from [79] and (vi) telomere PICh using BLM-deficient U2OS cells as PIChB (column Y) using data from [20].

#### Table 2 Overlap between TAILS and previously reported ALT modulators, Related to Figure 1.

#### Table 3 List of sgRNA sequences used in this study.
